# Convergent olfactory circuits for courtship in *Drosophila* revealed by *ds*-Tango

**DOI:** 10.1101/2024.10.23.619891

**Authors:** John D. Fisher, Anthony M. Crown, Altar Sorkaç, Sasha Martinez-Machado, Nathaniel J. Snell, Neel Vishwanath, Silas Monje, An Vo, Annie H. Wu, Rareș A. Moșneanu, Angel M. Okoro, Doruk Savaş, Bahati Nkera, Pablo Iturralde, Aastha Kumari, Cambria Chou-Freed, Griffin G. Hartmann, Mustafa Talay, Gilad Barnea

**Affiliations:** Department of Neuroscience, Brown University, Providence, RI, USA; Carney Institute for Brain Science, Brown University, Providence, RI, USA

**Keywords:** *ds*-Tango, disynaptic tracing, neural circuit, courtship, *Drosophila*, lateral horn

## Abstract

Animals exhibit sex-specific behaviors that are governed by sexually dimorphic circuits. One such behavior in male *Drosophila melanogaster*, courtship, is regulated by various sensory modalities, including olfaction. Here, we reveal how sexually dimorphic olfactory pathways in male flies converge at the third-order, onto lateral horn output neurons, to regulate courtship. To achieve this, we developed *ds*-Tango, a modified version of the monosynaptic tracing and manipulation tool *trans-*Tango. In *ds*-Tango, two distinct configurations of *trans-*Tango are positioned in series, thus providing selective genetic access not only to the monosynaptic partners of starter neurons but also to their disynaptic connections. Using *ds*-Tango, we identified a node of convergence for three sexually dimorphic olfactory pathways. Silencing this node results in deficits in sex recognition of potential partners. Our results identify lateral horn output neurons required for proper courtship behavior in male flies and establish *ds*-Tango as a tool for disynaptic circuit tracing.

## INTRODUCTION

Animals have evolved a variety of means to signal their suitability to potential mating partners. These signals relay information about the species, sex, and fitness of the animal via courtship rituals that consist of stereotyped sequences of behaviors, culminating in copulation. The courtship rituals of each species involve a specific array of sensory modalities that range from electroreception in electric fish^1^ to audition in bats^2^, from vision in birds of paradise^3^ to mechanosensation in spiders^4^.

In the fruit fly *Drosophila melanogaster*, courtship involves visual, auditory, tactile, gustatory, and olfactory signals^5–9^. Among the olfactory signals, volatile pheromones are used to correctly assess the suitability of a potential mate^6,10,11^, while environmental stimuli such as food odors affect the rate of copulation^12–14^. Since male and female fruit flies exhibit distinct mating behaviors^15^, they must respond differently to these olfactory stimuli. Indeed, several sexually dimorphic olfactory sensory pathways have been identified^16–18^.

Sexual dimorphism in the *Drosophila* nervous system is achieved by differential expression of sex-specific isoforms of the transcription factors fruitless (fru)^19,20^ and doublesex (dsx)^21^. Male-specific neurons express the male isoforms of these proteins (Fru^M^ and Dsx^M^). In males, the Fru^M^ isoform is present in olfactory sensory neurons (OSNs) that express the odorant receptors Or47b or Or67d, or the ionotropic receptor Ir84a^16–18^. Hence, these three populations of OSNs likely play a role in male-specific behaviors. Indeed, these receptors detect distinct aspects of the environment relevant for courtship. Or47b detects two pheromones that are produced by both sexes, methyl laurate^22^ and palmitoleic acid^23^, and the activation of Or47b-OSNs promotes copulation in males^24^. Or67d, on the other hand, binds to a male-specific pheromone, 11-*cis*-vaccenyl acetate (cVA)^25,26^, which is an aphrodisiac for females and a suppressor of mating in males^6^. By contrast, the ligands of Ir84a are not pheromones but rather food-derived odors such as phenylacetic acid and phenylacetaldehyde^12^. The activation of Ir84a-OSNs enhances male courtship^12^.

The presence of the Fru^M^ isoform in these three olfactory circuits and their roles in male mating behaviors suggest that they might interact. This interaction could occur at the first-order level of the circuits (OSNs), the second-order level (local interneurons (LNs) or olfactory projection neurons (OPNs)), or at a higher level. Although OSNs affect the electrical properties of each other through ephaptic coupling^27^; this is unlikely to happen among Or47b-, Or67d-, and Ir84a-OSNs as they reside in different sensilla^18,28^. In addition, while Or47b-driven courtship is suppressed by cVA, this effect requires synaptic release from the Or67d-OSNs^29^. Alternatively, the interaction between these three pathways could happen via LNs that innervate all three glomeruli. This hypothesis is unlikely since the activity of the Ir84a circuit is less prone to lateral inhibition from other olfactory circuits^30^. However, an important caveat of this study is that the inhibitory effects of the activation of the Or47b and Or67d circuits were not directly examined^30^. We, therefore, expect minimal contribution from the first- and second-order neurons in these circuits to the integration of olfactory courtship cues.

Thus, olfactory courtship cues may be integrated in higher-order neurons within these three circuits. Indeed, the OPNs in these circuits innervate a common region in the lateral horn (LH)^12,31^, a brain area that acts as a hub of integration for olfactory stimuli governing innate behaviors^31^. Despite extensive mapping of the various cell types that make up the LH^32,33^, our understanding of how neuronal activity in the LH influences innate behaviors is limited. Individual sexually dimorphic neurons in the LH that respond to cVA have been identified^34–36^, but no such population is known to receive inputs from the other olfactory circuits affecting courtship. Further, while cVA negatively regulates courtship^6^, the other Fru^M^+ circuits positively affect the courtship drive^12,24^. Thus, there remains a significant knowledge gap as to how olfactory information is routed through the brain to ultimately elicit male courtship behavior. We, therefore, sought to identify a potential node in the LH that translates neuronal activity in courtship-regulating olfactory circuits into the appropriate sexual behaviors.

To achieve this, we comprehensively mapped the Fru^M^+ olfactory pathways starting from the OSNs through three layers of neural circuitry to the LH. Given that the published fly connectomes are from female brains^37–39^, we could not trace these pathways in the male brain. We, therefore, developed a new version of *trans*-Tango^40^ that selectively labels neurons across two consecutive synapses downstream from a starting population. We termed this new method *ds*-Tango for disynaptic Tango. Using *ds*-Tango, we revealed that Or47b, Or67d, and Ir84a pathways converge onto a group of LH neurons that are third-order to all three circuits. Silencing of these neurons lead to increased male-to-male courtship. Our study identifies a new circuit node important for proper copulation behavior in *Drosophila melanogaster*.

## RESULTS

### Tracing second-order projections within the sexually dimorphic olfactory circuits

The antennal lobe (AL) is the first olfactory information processing center in the fly brain. In the AL, OSNs synapse onto OPNs that relay the olfactory information to higher-order brain regions, such as the mushroom body (MB) and the LH^41^. The MB is implicated in associative learning^42^ whereas the LH mostly mediates innate behaviors, including courtship^43^. To examine whether the three Fru^M^+ olfactory circuits converge on a common node in the LH, we mapped the second-order neurons within these circuits via *trans*-Tango, a genetically encoded circuit tracing method that works monosynaptically in the anterograde direction^40^ (Figure 1). In all three cases, we observed strong post-synaptic signal in the LNs, as well as the characteristic projections of the OPNs to the MB and LH (Figure 1). The LH is subdivided into discrete regions for different classes of odorants, with food related odors being represented in the dorsal LH, and pheromone odors being represented in the ventral LH^31,44^. Accordingly, the post-synaptic signal in both Or67d and Or47b circuits was localized strictly in the ventral LH (Figure 1A and 1B). By contrast, the OPNs of the Ir84a circuit innervated both the ventral and dorsal compartments (Figure 1C), consistent with the ligands for Ir84a being courtship-promoting food odors^12^. Thus, the three Fru^M^+ olfactory circuits may converge onto a common node in the ventral LH as this is a common projection target area. We, therefore, sought to map the circuits downstream of the OPNs to identify this putative node of convergence.

**Figure 1.**
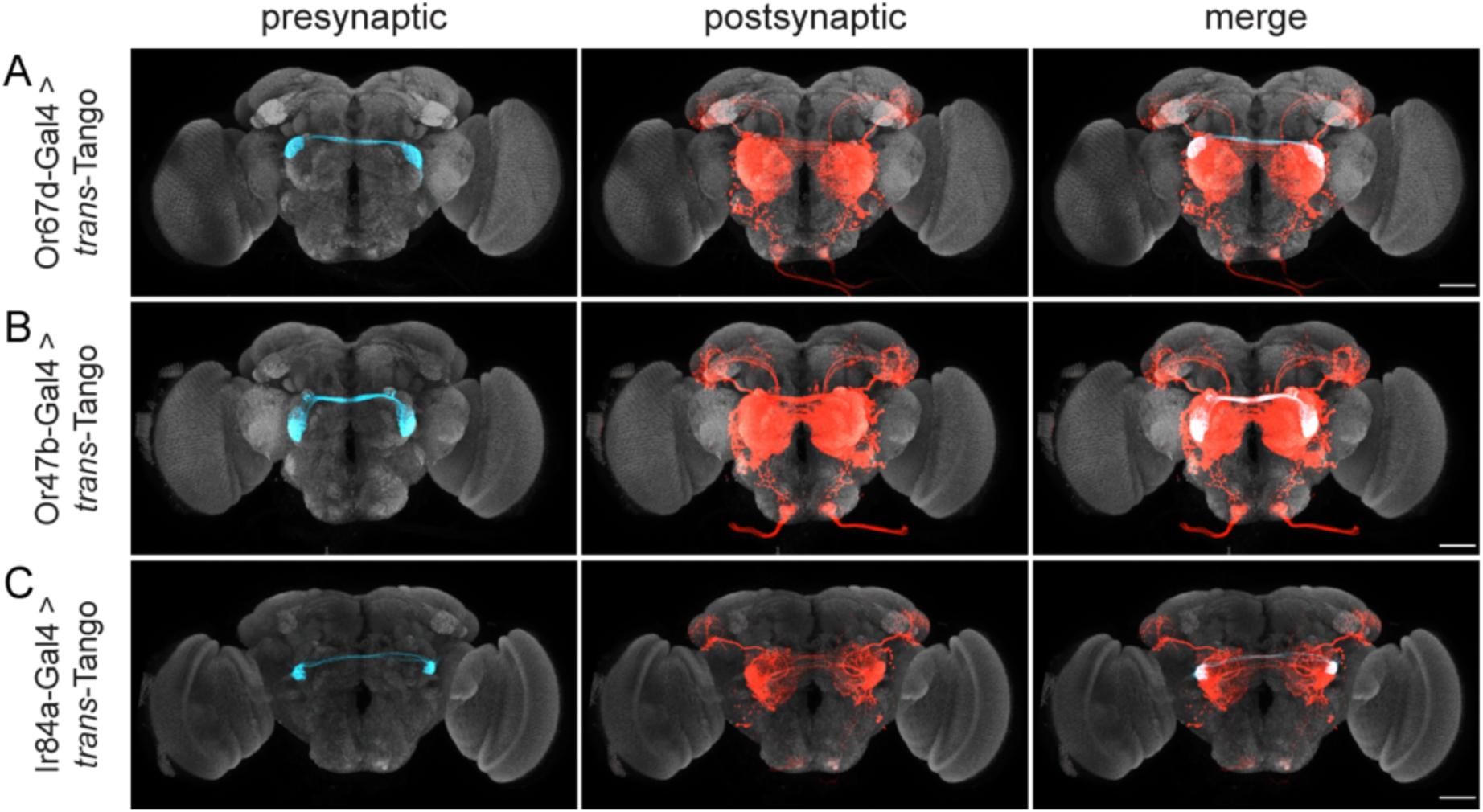
Tracing second-order projections within the sexually dimorphic olfactory circuits. *trans-*Tango is used to trace the projections of the three Fru^M^+ OSN populations: Or67d-OSNs (A), Or47b-OSNs (B), and Ir84a-OSNs (C). In the presynaptic channel (left column), only the OSNs are labeled (cyan). In the postsynaptic channel (center), both the LNs and PNs (red) are revealed. OPN axons target ventral LH in (A) and (B), and both ventral and dorsal LH in (C). Merge of both channels is shown in the rightmost column. Maximum intensity Z-stack projection of whole-mount brains are shown. Neuropil counterstain is shown in grey. Scale bars, 50μm.

### Design of *ds*-Tango

To map the third-order neurons in the Fru^M^+ olfactory circuits, we established a strategy for disynaptic anterograde tracing in the fly brain that we termed *ds*-Tango (For extended details on the design of *ds-*Tango, see supplemental text). In *ds*-Tango, two distinct versions of *trans-*Tango^40^ are configured in series, such that initiation of the first in the starter neurons leads to expression of the ligand of the second in the postsynaptic partners. This ligand, in turn, activates the second version of *trans-*Tango in third-order, disynaptic connections of the starter neurons. In this manner, *ds*-Tango enables selective genetic access into neurons mono- and disynaptic from the starter population.

The *ds*-Tango signaling pathway consists of five fusion proteins and three reporters (Figures 2A, S1, and S2). Three of the fusion proteins are expressed in all neurons: the human glucagon receptor tethered to a modified version of the transcription factor LexA^45^ via the cleavage site recognized by the tobacco etch virus N1a protease (hGCGR::TEVcs::LexA*), the human parathyroid hormone receptor tethered to the transcription factor QF by the same cleavage site (hPTHR::TEVcs::QF), and the human β-Arrestin2 fused to the N1a protease from the tobacco etch virus (hArr::TEV). The panneuronal expression of these components endows all neurons with the ability to react to both *ds*-Tango ligands, rendering the method flexible and versatile. The two remaining fusion proteins are the ligand constructs that are conditionally expressed in subsets of neurons. The first, a fusion between human glucagon and *Drosophila* Neurexin1 (hGCG::dNRXN1), is expressed in the starter neurons via the Gal4/UAS binary system. In the second, a modified version of human parathyroid hormone is similarly fused to *Drosophila* Neurexin1 (hPTH::dNRXN1). This fusion protein is expressed under the control of LexAop. To selectively label the three layers of the circuit, three reporter proteins are expressed under the control of the different binary systems. The starter neurons (shown in cyan) are labeled by CD2 under the control of Gal4/UAS, the monosynaptic partners (shown in green) are labeled by GFP under the control of LexA/LexAop, and the nuclei of the disynaptic connections (shown in magenta) are labeled by DsRed under the control of QF/QUAS^46^ (Figure 2A).

**Figure 2.**
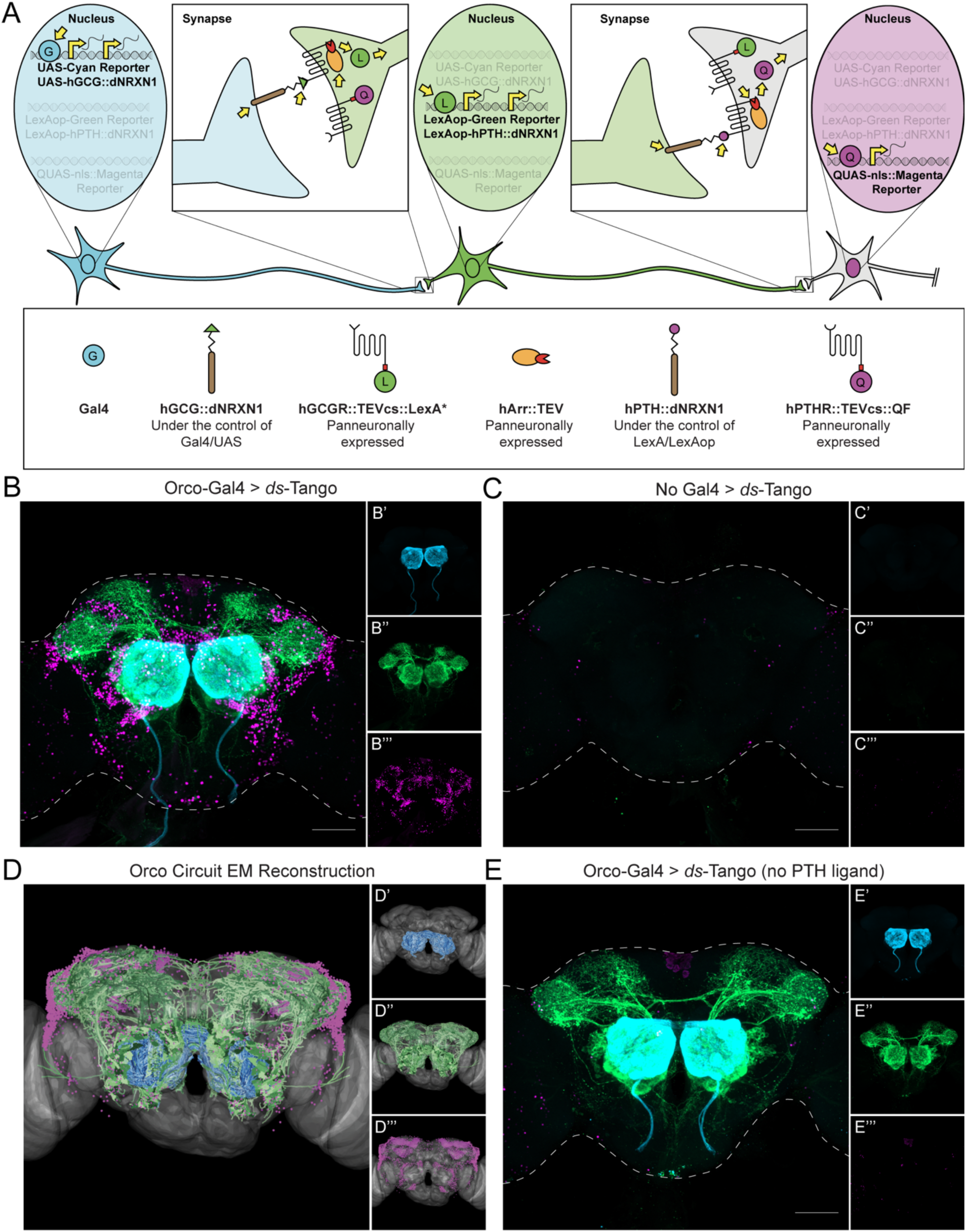
Design and implementation of *ds*-Tango in the olfactory system. (A) Schematic and components of *ds*-Tango. In flies carrying a Gal4 driver, the presynaptic reporter (cyan) and the GCG ligand (hGCG::dNRXN1) are expressed in the starter neurons. The GCG ligand localizes to the presynaptic sites of the starter neurons and activates the GCGR Tango fusion (hGCGR::TEVcs::LexA*) across the synapse on the monosynaptic partners. Upon activation of the GCGR, the hArr::TEV fusion protein is recruited to it, TEV cleaves its recognition site (TEVcs) releasing LexA*. LexA* then translocates to the nucleus and initiates the expression of the PTH ligand (hPTH::dNRXN1) and of the monosynaptic reporter (green). The PTH ligand localizes to the presynaptic sites of the monosynaptic partners and activates the PTHR Tango fusion (hPTHR::TEVcs::QF) across the synapse on the disynaptic connections. Upon activation of the PTHR, the hArr::TEV fusion protein is recruited to it, TEV cleaves its recognition site (TEVcs), releasing QF. QF then translocates to the nucleus and initiates the expression of the nuclear disynaptic reporter (magenta). The various steps in the process are indicated by yellow arrows. (B) Driving *ds*-Tango in the peripheral olfactory system using *Orco*-Gal4 labels OSNs (cyan, shown in B’), their monosynaptic partners LNs and OPNs (green, shown in B’’), and the nuclei of their disynaptic connections (magenta, shown in B’’’). (C) A brain of a control fly bearing the *ds*-Tango components, but no Gal4 driver exhibits no background neurons in the presynaptic channel (cyan, shown in C’), virtually no background neurons in the monosynaptic channel (green, shown in C’’), and the nuclei of a few background neurons in the disynaptic channel (magenta, shown in C’’’). (D) EM reconstruction of the Orco circuit projected on a template brain (gray) reveals OSNs (cyan, shown in D’), LNs and OPNs (green, shown in D’’), and the nuclei of third-order olfactory neurons (magenta, shown in D’’’). (E) A brain of a control fly bearing *Orco*-Gal4 and a version of *ds*-Tango lacking the PTH ligand exhibits labeling in OSNs (cyan, shown in E’), LNs and OPNs (green, shown in E’’), but no staining of disynaptic partners except for the nuclei of a few background neurons (magenta, shown in E’’’). Maximum intensity Z-stack projection of whole-mount brains are shown in B, D, and E. Dashed lines in B, D, and E depict the approximate outline of the fly brains. Scale bars = 50 μm.

The multilevel labeling scheme is initiated by Gal4 that, when expressed in a group of starter neurons, triggers the disynaptic *ds*-Tango signaling cascade (Figure 2A). In *ds*-Tango flies that do not carry a Gal4 driver, the GCG ligand is not expressed, and the *ds*-Tango cascade is not triggered. Thus, in all neurons, both QF and LexA* remain fused to their respective receptors and sequestered to the cell membrane, and none of the reporters are expressed. By contrast, in *ds*-Tango flies carrying a Gal4 driver, the CD2 reporter and the GCG ligand are expressed in all starter neurons. In these neurons, the GCG ligand construct targets to the presynaptic membrane, selectively initiating the GCGR::TEVcs::LexA* signaling cascade in their monosynaptic partners (Figure 2A). In the monosynaptic partners, the Arr::TEV fusion is recruited to the activated GCGR fusion, leading to cleavage of the TEVcs and releasing the transcription factor from the receptor. LexA* is now free to translocate to the nucleus where it initiates the expression of GFP and the PTH ligand construct. The PTH ligand construct is then targeted down the axon to the presynaptic membrane of the monosynaptic partners, allowing the ligand to bind its receptor on the disynaptic connections. This, in turn, initiates the PTHR::TEVcs::QF Tango signaling cascade in the disynaptic connections, culminating in translocation of QF to the nucleus and subsequent expression of nuclear DsRed. We used a nuclearly localized reporter as the final output for *ds*-Tango instead of a membrane-targeted reporter because the vast number of intermingled neurites labeled by the latter complicated analysis and troubleshooting. The modularity of *ds*-Tango enables one to readily replace the nuclearly localized reporter with other reporters, as shown below.

### Implementation of *ds*-Tango in the olfactory circuits

To validate *ds*-Tango, we initiated disynaptic tracing from OSNs to visualize the expected first-, second-, and third-order neurons in the olfactory circuits. To this end, we initiated *ds*-Tango using *orco*-Gal4, a driver that expresses in most OSNs (Figure 2B). We observed the stereotypical innervation patterns in the AL, and the characteristic OPN projections to the MB, LH, and subesophageal zone (SEZ). This confirmed that the *ds*-Tango signal in the second-order neurons recapitulates previously published *trans*-Tango tracing experiments^40^ and connectome results^47^. We also observed canonical third-order nuclei labeling around the AL, MB, and LH, consistent with the known anatomy of the olfactory circuits. In addition, we revealed third-order nuclei surrounding the SEZ, the primary taste center in the fly brain (Figure 2B), likely due to the projections from the AL to the SEZ^40^. To confirm that the *ds*-Tango labeling that we observed is Gal4-dependent, we examined *ds*-Tango flies lacking a Gal4 driver. Indeed, no first- or second-order background noise was present in these flies. We did, however, observe few third-order background neurons that were mostly restricted to the optic lobes (Figure 2C).

To further validate that the neurons revealed by *ds*-Tango are *bona fide* third-order olfactory neurons, we simulated a disynaptic tracing experiment using the *Drosophila* EM hemibrain dataset^37^. In our EM reconstructions of the olfactory circuit, we observed similar first-, second-, and third-order signal as we did in our olfactory *ds*-Tango experiment (Figure 2D). To confirm that the GCG fusion does not activate PTHR and that the third-order signal is PTH-dependent, we generated flies bearing all the *ds*-Tango components except for the PTH ligand fusion (Figure S2B). As expected, when we initiated this deficient configuration of *ds*-Tango with *orco*-Gal4, we observed similar first- and second-order labeling of neurons as with *ds*-Tango, but without labeling of third-order neurons (Figure 2E). These experiments confirmed the successful implementation of *ds*-Tango for selective labeling of neurons within the first three layers of the olfactory circuit.

### Specificity of *ds*-Tango

To examine the specificity of *ds*-Tango, we decided to initiate it from Or67d- and Or42a-OSNs, two populations we expect to have predominantly non-overlapping monosynaptic and disynaptic connections. Or67d is a pheromone receptor, and Or42a is broadly tuned to food odors^48^. Pheromones and food odors are represented in different compartments of the LH^31^. We, hence, sought to test whether *ds*-Tango would label distinct populations of LH neurons when initiated from these two OSN populations. Initially, we simulated *ds*-Tango experiments from the OSNs targeting the DA1 glomerulus (Or67d-OSNs) and OSNs targeting the VM7 glomerulus (Or42a-OSNs) using the *Drosophila* EM hemibrain dataset^37^ (Figures 3A and 3D). The EM reconstruction of the Or67d circuit revealed second-order projections to the ventral part of the LH and third-order neurons surrounding the LH (Figure 3A). Importantly, we noticed few to no cell bodies of the disynaptic connections surrounding the calyx of the MB. By contrast, the EM reconstruction of the Or42a circuit revealed second-order projections to the dorsal part of the LH, dense innervation of the MB, and numerous cell bodies of the disynaptic connections surrounding the calyx of the MB (Figure 3D). These differences serve as important benchmarks to evaluate the specificity of *ds*-Tango.

**Figure 3.**
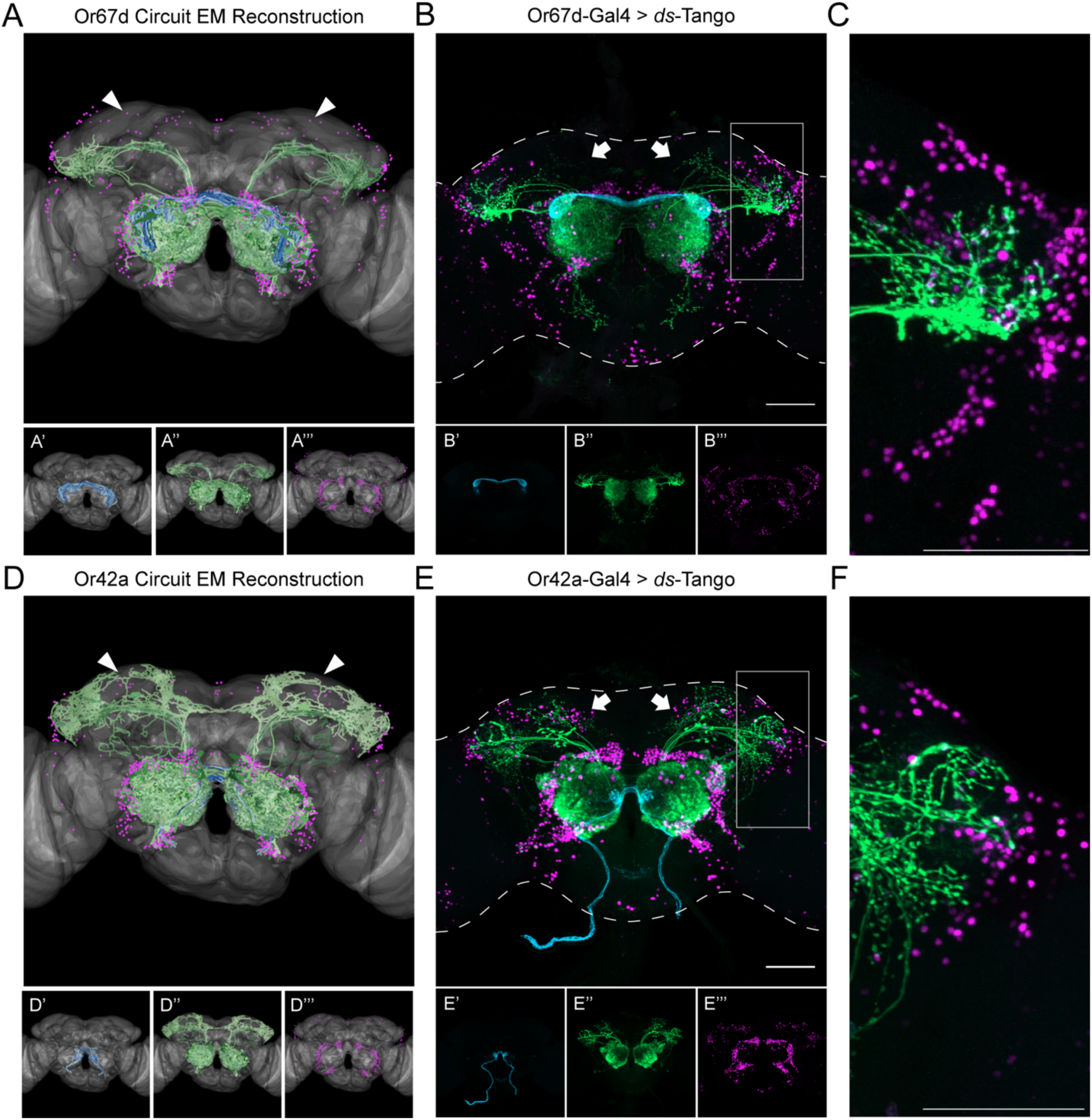
Specificity of *ds*-Tango. (A) Or67d Circuit EM reconstruction projected on a template brain (gray) reveals the OSNs (cyan, shown in A’), LNs and OPNs (green, shown in A’’), and the nuclei of third-order neurons (magenta, shown in A’’’). (B) Driving *ds*-Tango using *Or67d*-Gal4 labels Or67d-OSNs (cyan, shown in B’), their monosynaptic partner LNs and OPNs (green, shown in B’’), and the nuclei of their disynaptic connections (magenta, shown in B’’’). (C) Higher magnification image of the gray inset in (B), depicting the lateral horn. (D) Or42a Circuit EM reconstruction projected on a template brain (gray) reveals the OSNs (cyan, shown in D’), LNs and OPNs (green, shown in D’’), and the nuclei of third-order neurons (magenta, shown in D’’’). (E) Driving *ds*-Tango using *Or42a*-Gal4 labels Or42a-OSNs (cyan, shown in E’), their monosynaptic partners LNs and OPNs (green, shown in E’’), and the nuclei of their disynaptic connections (magenta, shown in E’’’). (F) Higher magnification image of the gray inset in (E), depicting the lateral horn. Arrowheads in A and D highlight the absence and presence of third-order Kenyon cells in the MB, respectively. Arrows in B and E highlight the absence and presence of third-order Kenyon cells in the MB, respectively. Maximum intensity Z-stack projection of whole-mount brains are shown in B and E. Dashed lines in B and E depict the approximate outline of the fly brains. Scale bars = 50 μm.

We then initiated *ds-*Tango from Or67d-OSNs (Figure 3B and 3C) and Or42a-OSNs (Figures 3E and 3F) and observed labeling in anatomically distinct populations of monosynaptic partners and disynaptic connections. Notably, initiation of *ds*-Tango from Or67d-OSNs identified second-order OPNs that densely innervate the ventral part of the LH as well as previously described non-canonical PNs that target the SEZ^40^ (Figure 3B and 3C). It is worth noting that our EM simulations of the Or67d circuit did not reveal second-order PNs targeting the SEZ, or third-order neurons surrounding the SEZ, because this brain area has not yet been traced and thus is not included in the EM hemibrain dataset^37^. By contrast, initiating *ds*-Tango from Or42a-OSNs revealed second-order PNs that target predominantly the calyx of the MB and the dorsal LH (Figures 3E and 3F). Notably, the projection to the SEZ observed when driving *ds*-Tango from Or67d-OSNs is absent when *ds*-Tango is driven from Or42a-OSNs. Further, we observed more third-order neurons surrounding the LH in the Or67d circuit than in the Or42a circuit (Figures 3C and 3F). By contrast, *ds*-Tango labeled more cell bodies of disynaptic connections surrounding the MB, when initiated from Or42a-OSNs (Figures 3B and 3E). Thus, the differences between the Or67d and Or42a circuits predicted by the EM reconstruction were successfully recapitulated by *ds*-Tango in both monosynaptic partners and disynaptic connections. These results demonstrate the accuracy and specificity of *ds*-Tango.

While troubleshooting *ds*-Tango, we identified several variables that affect the signal, including the rearing temperature and age of dissected flies. We observed an inverse relationship between temperature and the signal in monosynaptic partners and disynaptic connections (Figure S3). We noticed an optimal signal-to-noise ratio for standard tracing at 21°C, but rearing flies at either higher or lower temperatures can also provide benefits in some experimental designs. Further, we observed a direct relationship between the age of flies and the third-order *ds*-Tango signal (Figure S4). Therefore, users should optimize the protocol of their *ds*-Tango experiments according to their driver lines and circuits of interest. In this study, we crossed and raised flies at 21°C and dissected them at 3-4 weeks of age, unless otherwise noted. Using these parameters, we noticed remarkably consistent *ds*-Tango signal in the brains of different flies when driving the system with *Or67d*-Gal4 (Figure S5). On average, we observed 39.75±7.21 monosynaptic partners and 533±73.70 disynaptic connections per hemisphere across eight brains.

These variables, and others, can impact the *ds*-Tango signal, and thus impact the interpretation of the experimental results. Bias can be introduced in both EM-simulated *ds*-Tango reconstructions (via synaptic thresholds) and *ds*-Tango tracing (via experimental parameters such as age or temperature). Therefore, we sought other ways to validate that the *ds*-Tango signal we observe accurately reflects the biological ground truth.

### *ds*-Tango reveals sexual dimorphism in the lateral horn in the Or67d circuit

To further validate the specificity of *ds*-Tango in revealing differences between various neural pathways, we deployed it to explore sexually dimorphic circuits. Third order neurons in the Or67d circuit exhibit sexual dimorphism. While the Fru^M^+ lateral cluster neurons in the LH (LC1 and LC2) are part of the Or67d circuit in both sexes, the Fru^M^+ dorsal cluster neurons (DC1 and DC2) are postsynaptic to OPNs in males but not in females^34,35^. We wished to study whether *ds*-Tango is sufficiently specific to reveal this sexual dimorphism. To this end, we reasoned that a membrane bound reporter would enable easier identification of neurons, and thus, instead of the nuclearly localized one, we implemented a new reporter for the disynaptic connections, QUAS-mtdTomato^49^. In the absence of a Gal4 driver, this reporter exhibits little to no background in the central brain (Figures S6A and S6B). By contrast, when we initiated *ds*-Tango with this mtdTomato reporter from Or67d-OSNs, we observed dense disynaptic signal in the ventral portion of the LH in both males and females (Figures 4A and 4D). We also observed multiple distinct clusters of disynaptic neurons surrounding the LH in both sexes. Importantly, *ds*-Tango revealed sexual dimorphism in both the monosynaptic partners and disynaptic connections. We observed disynaptic LC1 and LC2 neurons in both sexes as expected^34,35^ (Figures 4A and 4D). By contrast, we noticed that the DC1 and DC2 neurons were strongly labeled in males; yet they were absent, or weakly labeled, in females (Figures 4A and 4D). To further confirm the identification of these neurons, we used an antibody against Fru^M^ alongside *ds*-Tango. We observed that among the disynaptic connections, not all neurons in the lateral clusters were Fru^M^+. By contrast, we did not observe co-labeling of Fru^M^ within the dorsal clusters (Figure S7). This result contradicts previous reports that observed Fru^M^ immunoreactivity in dorsal cluster neurons^35^. A potential explanation for this discrepancy is that we analyzed flies at an old age (three to four-week-old). To resolve this discrepancy, we restricted the expression of the disynaptic reporter via an intersectional genetic approach using a FLP recombinase expressed from the *fru* locus (*fru*-FLP)^50^ together with our FLP-dependent reporter. Our results with this approach further confirmed that while *ds*-Tango reveals lateral cluster neurons to be disynaptic connections of Or67d-OSNs in both sexes, the dorsal cluster neurons appear to be part of the Or67d circuit in males only (Figures 4B, 4C, 4E, and 4F). Importantly, this intersectional approach yielded little to no background signal in the absence of a Gal4 driver (Figures S6C and S6D). Altogether, these experiments validate that *ds*-Tango is sufficiently specific to reveal differences in sexually dimorphic circuits.

**Figure 4.**
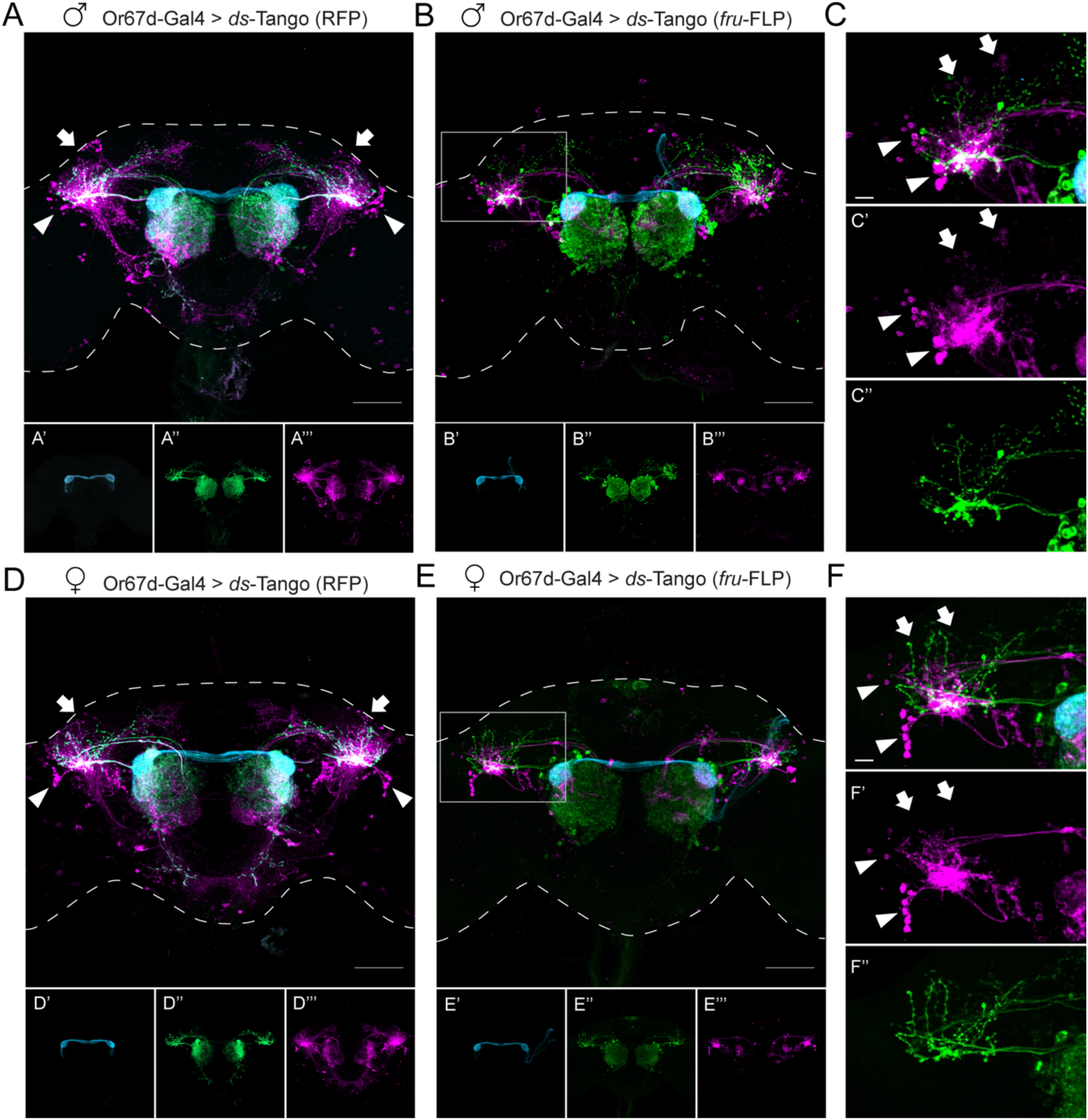
*ds*-Tango reveals sexual dimorphism in the lateral horn in the Or67d circuit. (A) Driving *ds*-Tango with *Or67d*-Gal4 in males labels Or67d-OSNs (cyan, shown in A’), their monosynaptic partner LNs and OPNs (green, shown in A’’), and their disynaptic connections (magenta, shown in A’’’). Arrows indicate the dorsal cluster neurons; arrowheads indicate the lateral cluster neurons. (B) Driving *ds*-Tango with *Or67d*-Gal4 and genetically restricting disynaptic reporter expression to *fru*-FLP+ neurons in males labels Or67d-OSNs (cyan, shown in B’), their monosynaptic partner LNs and OPNs (green, shown in B’’), and their *fru*-FLP+ disynaptic connections (magenta, shown in B’’’). (C) A higher magnification image of the gray inset in (B) highlighting the left LH reveals PNs targeting the ventral region of the LH (green, shown in C’’), overlapping with the neurites of *fru*-FLP+ disynaptic connections (magenta, shown in C’). Note the presence of lateral cluster neurons (arrowheads) and the dorsal cluster neurons (arrows) in the male brain. (D) Driving *ds*-Tango with *Or67d*-Gal4 in females labels Or67d-OSNs (cyan, shown in D’), their monosynaptic partner LNs and OPNs (green, shown in D’’), and their disynaptic connections (magenta, shown in D’’’). Arrowheads indicate the lateral cluster neurons; arrows indicate the location of dorsal cluster neurons. Note the presence of lateral cluster neurons in both the male (arrowheads in A) and female (arrowheads in D) brains and the prominent dorsal cluster neurons in the male (arrows in A) brain that are less prominent or absent in the female (arrows in D) brain. (E) Driving *ds*-Tango with *Or67d*-Gal4 and genetically restricting disynaptic reporter expression to *fru*-FLP+ neurons in females labels Or67d-OSNs (cyan, shown in E’), their monosynaptic partners LNs and OPNs (green, shown in E’’), and their *fru*-FLP+ disynaptic connections (magenta, shown in E’’’). (F) A higher magnification image of the gray inset in (E) showing the left LH reveals PNs targeting the ventral region of the LH (green, shown in F’’), overlapping with the neurites of *fru*-FLP+ disynaptic connections (magenta, shown in F’). Note the presence of lateral cluster neurons (arrowheads) and the absence of dorsal cluster neurons (arrows) in the female brain. Maximum intensity Z-stack projection of whole-mount brains are shown in A, B, D, and E. Dashed lines in A, B, D, and E depict the approximate outline of the fly brains. Scale bars, 50 μm.

### A node of convergence for courtship-regulating olfactory circuits

Having established *ds*-Tango as a specific tool for tracing disynaptic connections within a circuit, we turned to identifying nodes of convergence for courtship-regulating olfactory circuits. To this end, we sought to map the circuitry downstream from the two remaining Fru^M^+ OSN populations. The use of the *ds*-Tango strategy allows us to identify common targets of the Fru^M^+ OSNs in the LH. We first implemented *ds*-Tango to reveal all the third-order neurons in the Or47b and Ir84a circuits. We drove the configuration of *ds*-Tango in which the disynaptic reporter is mtdTomato from Or47b-OSNs and Ir84a-OSNs. We found that in each circuit the *ds*-Tango signal in the monosynaptic partners (Figures 5A and 5C) recapitulates the signal observed with *trans*-Tango (Figures 1B and 1C). Driving *ds*-Tango from Or47b-OSNs revealed innervation of the ventral compartment of the LH by the OPNs (Figure 5A). By contrast, driving *ds-*Tango from Ir84a-OSNs revealed OPNs that innervated the dorsal and ventral compartments of the LH (Figure 5C). Amongst the third-order neurons in both circuits, we observed ventral projections from the LH to the anterior ventrolateral protocerebrum (AVLP). However, when driving *ds*-Tango from Or47b-, or Ir84a-OSNs, the signal in the disynaptic connections was dense and hard to parse. We thus concluded that to identify a common node among the Fru^M^+ olfactory circuits, a sparser *ds*-Tango signal in the third-order neurons would be necessary.

**Figure 5.**
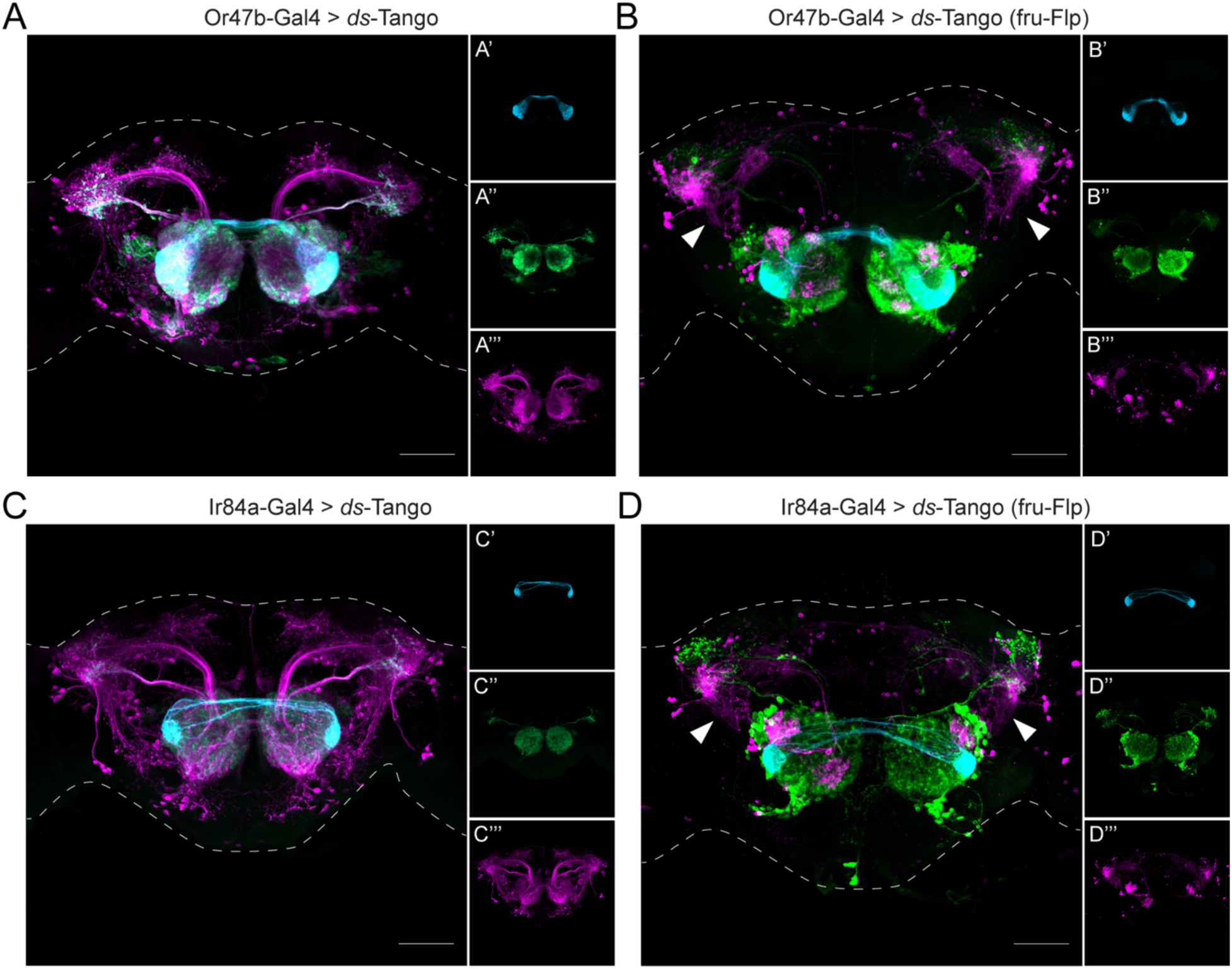
A node of convergence for courtship-regulating olfactory circuits. (A) Driving *ds*-Tango with *Or47b*-Gal4 in males labels Or47b-OSNs (cyan, shown in A’), their monosynaptic partner LNs and OPNs (green, shown in A’’), and their disynaptic connections (magenta, shown in A’’’). (B) Driving *ds*-Tango with *Or47b*-Gal4 and genetically restricting disynaptic reporter expression to *fru*-FLP+ neurons in males labels Or47b-OSNs (cyan, shown in B’), their monosynaptic partner LNs and OPNs (green, shown in B’’), and their *fru*-FLP+ disynaptic connections (magenta, shown in B’’’). Note the presence of ventral projections from the LH to the AVLP in the disynaptic connections characteristic of AV2b1/b2 neurons (arrowheads). (C) Driving *ds*-Tango with *Ir84a*-Gal4 in males labels Ir84a-OSNs (cyan, shown in C’), their monosynaptic partners LNs and OPNs (green, shown in C’’), and their disynaptic connections (magenta, shown in C’’’). (D) Driving *ds*-Tango with *Ir84a*-Gal4 and genetically restricting disynaptic reporter expression to *fru*-FLP+ neurons in males labels Ir84a-OSNs (cyan, shown in D’), their monosynaptic partners LNs and OPNs (green, shown in D’’), and their *fru*-FLP+ disynaptic connections (magenta, shown in D’’’). Note the presence of ventral projections from the LH to the AVLP in the disynaptic connections characteristic of AV2b1/b2 neurons (arrowheads). Maximum intensity Z-stack projection of whole-mount brains are shown. Dashed lines depict the approximate outline of the fly brains. Scale bars, 50 μm.

We reasoned that a hypothetical LH neuron that would integrate information from the three Fru^M^+ olfactory circuits would also express Fru^M^. We, therefore, restricted the disynaptic *ds*-Tango signal using *fru*-FLP together with the FLP-dependent configuration of *ds*-Tango (Figure S2D). Driving this configuration of *ds*-Tango from Or47b- or Ir84a-OSNs labeled smaller populations of disynaptic connections (Figures 5B and 5D) than in the analogous experiments using the configuration of *ds*-Tango that does not require FLP expression (Figures 5A and 5C). When we compared the fru-FLP+ third-order neurons in the Or67d (Figure 4B), Or47b (Figure 5B), and Ir84a (Figure 5D) circuits, we noticed a common tract that projects ventrally from the LH into the AVLP. This projection is characteristic of AV2b1/b2 neurons^33^, a subset of the LC2 cluster^34^. Due to the distinct morphology of these neurons that resemble the meandering river Maiandros, we named them the Maiandros neurons. We reasoned that the Maiandros neurons could serve as a node of convergence that relays courtship-relevant olfactory cues to downstream processing centers to regulate courtship behaviors.

### Silencing the Maiandros neurons leads to deficits in mate discrimination in males

We next sought to characterize the Maiandros neurons through the use of a specific genetic driver line. We screened through the expression patterns of split-Gal4 drivers in a library that targets various cell types of the LH^33^. One such driver, LH2088-Gal4^33^, provides weak access to the Maiandros neurons with only faint background in some of the Kenyon cells of the MB (Figure 6A). Using this driver, we sought to determine the polarity of the Maiandros neurons in males by expressing markers for presynaptic sites (synaptotagmin::GFP^51^) and dendrites (DenMark^52^). We reasoned that if the Maiandros neurons indeed relay information from the LH to downstream circuits, their dendrites should be localized in the LH, and their axons should be localized in the AVLP. Visualizing the neurite positions of the Maiandros neurons revealed that they have mixed axons and dendrites in the LH but only axons in the AVLP (Figure 6B). Because the neurites in the AVLP were strictly axonal, we conclude that the Maiandros neurons in males relay information from the LH to the AVLP as predicted for females^53,54^, indicating that they are indeed LH output neurons. In females, these neurons were determined to be both GABAergic and cholinergic through neurotransmitter staining^33^, yet only cholinergic by electron microscopy reconstructions^53^. To identify the neurotransmitters used by these neurons in males, we examined the expression of choline acetyltransferase (ChAT - cholinergic) and glutamic acid decarboxylase (GAD - GABAergic) drivers in these neurons. We observed that only the ChAT driver expressed in these neurons (Figure S8), indicating that in males these neurons are likely cholinergic.

**Figure 6.**
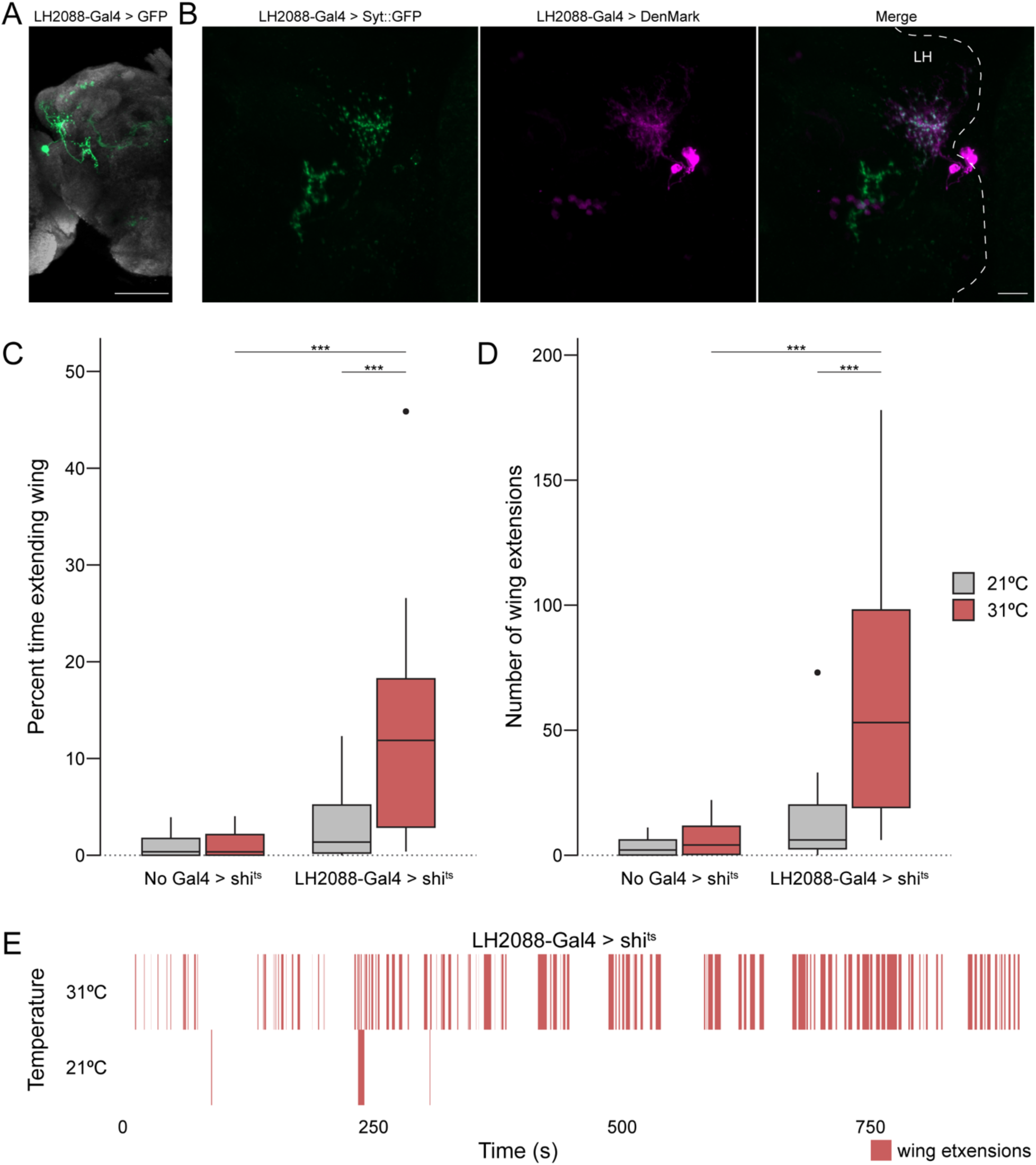
Anatomical and functional analysis of the Maiandros neurons. (A) Expression pattern of the LH2088-Gal4 driver line. LH2088-Gal4 drives UAS-GFP (green). The Maiandros neurons are clearly labeled along with minimal background in the MB. Neuropil counterstain is shown in grey. (B) Locations of the Maiandros neuron axons and dendrites. LH2088-Gal4 drives synaptic vesicle marker Syt::GFP (green) and dendritic marker DenMark (magenta). Axons and dendrites are intermixed in the LH while ventral neurites in the AVLP are primarily axons. Dashed lines indicate the outline of the central brain. LH indicates lateral horn. (C) Quantification of unilateral wing extension as indicator of attempted courtship when each experimental male fly is paired with a single naïve WT male. Boxplot depicting the percent of total trial time (15 min) that experimental male flies extended one wing during thermogenetic silencing using shi^ts^. Silencing of the Maiandros neurons in male flies tested at the restrictive temperature of 31°C results in increased time spent with one wing extended relative to genetic controls and to experimental flies tested at the permissive temperature of 21°C. (D) Quantification of unilateral wing extension as indicator of attempted courtship when each experimental male fly is paired with a single naïve WT male. Boxplot depicting the total number of unilateral wing extensions exhibited by experimental male flies during thermogenetic silencing using shi^ts^. Silencing of the Maiandros neurons in male flies tested at the restrictive temperature of 31°C results in increases in total number of wing extensions relative to genetic controls and to experimental flies tested at the permissive temperature of 21°C. (E) Rasterplot depicting bouts of unilateral wing extensions over time for representative trials of male flies bearing the LH2088-Gal4 and UAS-shi^ts^ alleles in the thermogenetic silencing assay at the permissive temperature of 21°C and the restrictive temperature of 31°C. Unilateral wing extensions are sparse at 21°C but abundant throughout the 31°C trial. Maximum intensity Z-stack projection of whole-mount brains are shown in (A) and (B). Scale bars, 50μm in (A) and 10μm in (B).

Because the Maiandros neurons receive synaptic inputs from the courtship-regulating olfactory circuits, we hypothesized that their activity would be necessary for proper control of courtship behavior in males. Under natural conditions, flies only exhibit minimal levels of intermale courtship due to pheromonal cues, such as cVA, that suppress courtship behaviors^6,55^. Thus, we tested whether silencing the Maiandros neurons would lead to intermale courtship. To achieve this, we used a temperature sensitive Dynamin allele (shi^ts^)^56^ that reversibly blocks neurotransmission at temperatures above 28°C. In each trial, we paired a single experimental male with a single naïve w^11^^18^ male in a circular behavior arena^57^ and used the unilateral wing extension behavior^15^ as a readout of attempted courtship. Predictably, control males that carried the shi^ts^ allele but no driver exhibited only low levels of courtship towards a w^11^^18^ male at both 21°C and 31°C (Figures 6C and 6D). Flies expressing shi^ts^ in the Maiandros neurons exhibited a slight, albeit insignificant, increase in courtship behavior at the permissive temperature of 21°C. At the restrictive temperature of 31°C, however, these flies exhibited dramatically increased levels of intermale courtship (Figures 6C and 6D). Experimental males typically pursued w^11^^18^ males throughout the duration of the 15-minute trial, indicating that the courtship pursuits persisted long after they would have presumably detected the pheromones on their targets (Figure 6E). Thus, silencing the Maiandros neurons elicits a prolonged arousal state that suppresses the effects of the anti-aphrodisiac pheromones present on the courtship targets. We interpret these data as evidence that activity in the Maiandros neurons is required for proper courtship behaviors in male flies. We conclude that the Maiandros neurons constitute a node of convergence for all three sexually dimorphic olfactory circuits and that their activity is required for proper identification of the sex of potential mates.

## DISCUSSION

Control of innate behaviors, such as courtship, requires hardwired circuits to integrate a diverse array of sensory cues that are detected by multiple sensory systems. Even within a sensory modality, cues from various sources may be needed to convey diverse pieces of the information necessary for the proper execution of a certain behavior. For example, courtship in *Drosophila melanogaster* is regulated by sensory cues that are detected by at least three olfactory sensory channels. These cues may originate from conspecifics (*e.g.*, the pheromones cVA and methyl laurate), or from the environment (*e.g.*, the food odor phenylacetaldehyde). Regardless of their origin, these cues can either promote or inhibit courtship. Hence, the brain must integrate these distinct, and at times opposing, olfactory cues into a cohesive behavior. This integration may take place at a node of convergence for these three olfactory channels. Therefore, gaining a deeper understanding of how the brain transforms olfactory cues into courtship behaviors requires detailed mapping of the points of interaction between these channels. Since the LH is known to mediate innate behaviors, including courtship, it is well positioned to serve as the locus of this node of convergence.

The LH receives projections from the olfactory system that segregate into compartments based on the associated behaviors for the respective odors. Food odors are represented in the dorsal LH, and pheromones in the ventral LH^31,58^. However, whether this pattern of segregation of the OPN inputs into the LH is manifested as convergence onto a common node of output remained unknown. To address this, we developed *ds*-Tango, a genetic tool for tracing the disynaptic connections of starter neurons. Using *ds*-Tango, we traced the disynaptic connections of the three Fru^M^+ OSN populations that detect courtship-regulating olfactory cues in males. Our analysis revealed the previously catalogued AV2b1/b2 neurons^33^, which we named the Maiandros neurons, as a common target receiving convergent inputs from the three courtship-regulating olfactory circuits. The anatomy of the Maiandros neurons indicates that they receive olfactory information from the OPNs projecting to the LH and relay this information to downstream targets in the AVLP. Thus, the AVLP is well positioned to inherit courtship-regulating olfactory signals from the LH and serve as a higher-order brain region for processing courtship-relevant sensory information into the appropriate behaviors. Consistent with this hypothesis, silencing the Maiandros neurons led to intermale courtship, likely due to increased arousal and impaired mate discrimination. The Maiandros neurons may, therefore, play a key role in processing courtship-relevant odor information to control arousal levels and decide whether to court or not to court. Since silencing of the Maiandros neurons in males leads to increases in intermale courtship, these neurons are likely inhibited by courtship promoting pheromones and food odors and activated by antiaphrodisiac pheromones. Indeed, in females these neurons have been described to receive inhibitory signals upon activation of the Or47b pathway^59^. Further work is required to determine how activity in the Maiandros neurons is affected in males by the three sexually dimorphic olfactory pathways. The Maiandros neurons may regulate levels of arousal in the male fly, perhaps through indirect inhibitory signals onto P1 neurons – a population that regulates arousal and the intensity of courtship behaviors^60–63^.

To identify third-order neurons within the olfactory circuits, we developed *ds*-Tango, a genetically encoded synthetic signaling pathway that allows differential genetic access into the monosynaptic partners and disynaptic connections of a given population of starter neurons. While we demonstrate the efficacy of *ds*-Tango by implementing it in the well-described *Drosophila* olfactory system, a major advantage of *ds-*Tango is its versatility. The signaling pathway that mediates *ds-*Tango labeling is expressed panneuronally. *ds-*Tango can, therefore, be deployed to achieve disynaptic tracing from any neuron for which genetic access is available via a Gal4 driver. While here the starter neurons are the first-order, sensory neurons of the olfactory circuits, *ds*-Tango can be initiated from any layer within any circuit. Additionally, the reporters that *ds*-Tango selectively expresses in the starter neurons, monosynaptic partners, and disynaptic connections are all genetically encoded. This design enables researchers to vary the types of effectors that are expressed in each layer of a circuit. Indeed, this modularity has enabled the implementation of various configurations of the precursor of *ds*-Tango, *trans-* Tango, for calcium imaging^46,64^ as well as optogenetic^64,65^ and chemogenetic^66^ manipulations. *ds-*Tango is, therefore, not limited to tracing experiments and can be extended to enable the recording or manipulation of neuronal activity via the extensive *Drosophila* genetic toolkit.

The success of a *ds-*Tango experiment, however, depends on the Gal4 driver line being used. Leaky or non-specific expression in a given Gal4 line will confound results by producing false positive *ds*-Tango signal in addition to *bona fide* monosynaptic partners and disynaptic connections. To minimize the false positive signal, we recommend selecting Gal4 driver lines that narrowly target the desired starting population, when possible. Importantly, all the alleles for the *ds-*Tango system can be carried by a single fly. Consequently, performing a *ds-*Tango experiment only requires researchers to cross flies bearing the Gal4 driver labeling their starter neurons of choice to flies carrying the *ds*-Tango alleles and analyze the progeny of this cross. This feature renders feasible the use of *ds-*Tango with intersectional genetic strategies such as split-Gal4s^67^. However, our attempts to trace the neurons downstream of the Maiandros neurons using *ds-*Tango were not successful, likely due to the low levels of expression induced by the LH2088 split-Gal4 driver. Therefore, obtaining a Gal4 driver line that is both sufficiently strong and specific to the population of interest is imperative when designing a *ds*-Tango experiment.

The recently completed connectome of a *Drosophila* brain^37^ and other EM volumes that are readily accessible via various databases, such as the Virtual Fly Brain^68^, have become the gold standard in profiling connectivity in the fly. Nevertheless, *ds*-Tango complements the use of these connectomic strategies in several ways. First, EM reconstruction of a connectome requires expertise and is costly, laborious, and time intensive. By contrast, *ds*-Tango is user-friendly and only requires basic maintenance of a fly line. Second, the information within each EM volume only represents a single fly of a single sex at a single timepoint raised in a single environment. Thus, the use of EM volumes does not allow for comparison between different group of individuals for the study of the effect of age, sex, environment, past experiences or genetic background. By contrast, the ease of use of *ds*-Tango and its inherent high throughput allow researchers to readily profile circuit connectivity across multiple individuals, sexes, or various other conditions. The utility of these features has been exemplified by the use of the other members of the Tango toolkit – *trans-*Tango and *retro*-Tango – to study the effects of environment^69^, sex^49^, and genetic background^70,71^ on brain wiring. Here, we demonstrate these features by deploying *ds-*Tango to map the courtship-regulating olfactory circuits in males – a study we would not have been able to conduct since the only currently available EM volumes are from female brains.

Mapping synaptic connections in the brain is an essential step towards understanding how sensory information is processed to elicit the appropriate behaviors to the environment. Our study introduces *ds*-Tango, a new tool that can be deployed to analyze various circuits in the fly brain. *ds*-Tango is the first transsynaptic labeling tool that provides selective genetic access to three layers within a given neural circuit and thus endows researchers with the capacity to differentially manipulate neurons within each layer. We demonstrate the utility of *ds-*Tango by implementing it in the olfactory circuits for courtship behaviors in the male. This study provides a step forward in our understanding of male courtship behavior in *Drosophila melanogaster* by unveiling a potential node of integration for courtship-regulating sensory cues. The underlying network structure we describe may generalize to other pathways in the LH, such as the ones underlying food seeking and aversion. Our experiments thus lay the groundwork for a deeper understanding of how the brain controls innate behaviors.

## METHODS

### Fly Strains

All *Drosophila melanogaster* lines used in this study were raised on cornmeal-agar media supplemented with the antifungal Tegosept. Flies were maintained in 70% humidity-controlled incubators with a 12-hour light/dark cycle. Flies were crossed and raised at a single temperature: 18°C, 21°C, or 25°C, unless otherwise stated. Temperature details for all experiments are available in Table S1. The publicly available fly lines used in this study are as follows: *trans-*Tango (BDSC #77123), *fru*-FLP^50^, *Or42a*-Gal4 (BDSC #9969), *Or47b-*Gal4 (BDSC #9983), *Ir84a-* Gal4 (BDSC #41750), *Or67d*-Gal4^6^, *Orco*-Gal4 (*Or83b*-Gal4; BDSC #23292), QUAS-FRT-STOP-FRT-mCD8::GFP (BDSC #30134), QUAS-nlsRFP^46^, w^11^^18^ (BDSC #5905), LH2088-Gal4 (BDSC #86635), UAS-GFP (pJFRC81-10xUAS-Syn21-myr::GFP-p10)^72^ (Pfeiffer, Truman, and Rubin 2012), UAS-Sytaptotagmin::GFP (BDSC #6926), UAS-DenMark (BDSC #33061), UAS-Shibire(ts)^56^, SS25111-Gal4 (BDSC #86796), UAS-CsChrimson (BDSC #55135) and QUAS-tdTomato-HA^49^. The transgenic *ds*-Tango fly line that was generated for this study includes four new fly lines: elav-Arr::TEV_nSyb-PTHR::TEVcs::QF_gypsy insulator(INS)_nSyb-GCGR::TEVcs::LexA* (inserted at attP40), UAS-CD2_ gypsy insulator(INS)_UAS-GCG::dNRX1 (inserted at attP2), LexAop-tdTomato_ gypsy insulator(INS)_LexAop-PTH::dNRX1 (inserted at VK00027) and LexAop-GF_gypsy insulator(INS)_LexAop-PTH::dNRX1 (inserted at VK00027). Three other lines were also generated for use in this study: LexAop-GFP (inserted at VK00027), and LexAop-tdTomatoLexAop-PTH::dNRX1 (inserted at VK00027). More detailed information about the genetic components of *ds*-Tango flies and other flies used in this study are available in Figures S1 and S2. In addition, genotypes for flies used in all experiments are listed in Table S1.

### Generation of Transgenic Fly Lines

All plasmids generated for this study were generated using standard molecular biology techniques: restriction digest of a vector plasmid, PCR amplification of DNA inserts with Gibson overhangs added to primers (NEB, Q5 High-Fidelity DNA Polymerase), and Gibson assembly of all components (NEB, NEBuilder HiFi DNA Assembly). Gibson assembly mix was transformed into competent cells (NEB, NEB Stable Competent *E. coli*), cells were spread on antibiotic resistant plates, colonies were screened and sequenced, and finally sent for injection and insertion into appropriate attP sites via PhiC31 integration (Rainbow Transgenic Flies Inc., Camarillo, CA). Detailed DNA sequence and integration site information and of injected plasmids is in Supplemental Figure and the Results section. Complete sequence map information of any constructs used in this study are available upon request. The vector backbone plasmid and the insert template plasmids used to generate flies in this study are listed below.

elav-Arr::TEV_nSyb-PTHR::TEVcs::QF_gypsy insulator(INS)_nSyb-GCGR::TEVcs::LexA* was generated using the *trans*-Tango plasmid^40^ as the vector backbone. The PTHR sequence was amplified from a synthesized DNA fragment containing human PTHR (GENEWIZ). The non-PTHR encoding sequences for the PTHR fusion protein were amplified from the *trans*-Tango plasmid^40^. The gypsy insulator sequence (INS) was amplified from the plasmid pJFRC10-10XUAS-IVS-mCD8::GFP, INS, 10XUAS-IVS-mCD8::GFP^72^. The nSyb-GCGR::TEVcs::LexA* sequence was partially amplified from the *trans*-Tango plasmid^40^. For the LexA* part, the LexADBD sequence was amplified from Addgene #46117 and and the QFAD* from #61310^45^.

UAS-CD2, INS, UAS-GCG::dNRX1 was generated using the plasmid pJFRC10-10XUAS-IVS-mCD8::GFP, INS, 10XUAS-IVS-mCD8::GFP^72^ as the vector backbone. The UAS-CD2 sequence was amplified from the plasmid UAS-CD2_QUAS-mtdTomato(3xHA) (Scaplen et al. 2021). The UAS-GCG::dNRX1 sequence was amplified from the plasmid *trans*-Tango MkII^73^.

LexAop-GFP was generated using the pJFRC19-13XLexAop2-IVS-myr::GFP plasmid^72^ as the vector backbone and the “GFP” sequence was amplified from the pJFRC206-10XUAS-FRT>STOP>FRT-myr::smGFP-V5 plasmid^74^.

LexAop-GFP, INS, LexAop-PTH::dNRX1 was generated using the LexAop-GFP plasmid (this study) as the vector backbone. The INS sequence was amplified from plasmid UAS-CD2, INS, UAS-GCG::dNRX1 (this study). The LexAop-PTH::dNRX1 sequence was amplified from the following plasmids: LexAop-GFP plasmid (this study) — for the LexAop sequence and the sv40pA after the PTH ligand coding region — and the UAS-CD2, INS, UAS-GCG::dNRX1 plasmid (this study) — for the non-PTH encoding part of the PTH ligand. The PTH ligand sequence was generated using overhangs from Gibson primers. The PTH ligand sequence encodes all but the first amino acid from the propeptide and the entire mature PTH peptide: SVKKRSVSEIQLMHNLGKHLNSMERVEWLRKKLQDVHNFVALGAPLAPRDAGS QRPRKKEDNVLVESHEKSLGEADKADVNVLTKAKSQ We found that this specific peptide sequence gave a higher signal-to-noise ratio than the entire prepro-PTH peptide sequence, just the mature PTH peptide, or any other versions of the PTH peptide we generated (data not shown).

LexAop-tdTomato, INS, LexAop-PTH::dNRNX1 was generated using the LexAop-GFP, INS, LexAop-PTH::dNRX1 (this study) as the vector backbone. The tdTomato(3XHA) insert sequence was amplified from the QUAS-tdTomato(3XHA) plasmid^49^.

#### Immunohistochemistry and Tissue Processing

The age of each dissected experimental fly in this study is listed in Table 3-S1. Immunohistochemistry was performed on dissected adult fly brains as previously described^40^, with some adaptations. Briefly, adult flies were cold anesthetized on ice and dissected in 0.5% PBST. All following steps were performed while brains were nutating. Dissected brains were then fixed for 30 minutes at room temperature (RT) in 4% PFA/0.01% PBST, washed 4 times — each wash 15 minutes at RT in 0.5% PBST, and blocked for 90 minutes at RT in 5% heat-inactivated donkey serum (diluted in 0.5% PBST). Then, brains were incubated in primary antibodies (diluted in the donkey blocking solution) for two overnights. Following primary antibody incubation, brains were once again washed 4 times — each wash 15 minutes at RT in 0.5% PBST. Next, brains were incubated in secondary antibodies (diluted in the donkey blocking solution) for two overnights. Finally, brains were once again washed 4 times — each wash 15 minutes at RT in 0.5% PBST — and placed in 0.5% PBST for subsequent DPX clearing. The Janelia FlyLight Clearing Protocol “DPX Mounting” (https://www.janelia.org/sites/default/files/Project%20Teams/Fly%20Light/FL%20Protoc Pr%20-%20DPX%20Mounting%202020-03-06.pdf) was followed exactly with a few modifications. First, we used Fisher Scientific coverslips for mounting brains (Cover glass 22×22 mm Square No. 1, Fisher Scientific #12-542B), as was done in the original DPX Mounting protocol, not Corning brand. As the protocol authors noted in the updated protocol, we also observed inconsistent and incomplete PLL coating using Fisher brand coverslips. For this reason, we made our second modification to the protocol, by increased the PLL coating of coverslips step to 5-10 seconds at RT. An abbreviated version of the DPX Mounting protocol is below including a few modifications.

Following wash of secondary antibodies, brains were postfixed for 4 hours at RT in 4% PFA/0.01% PBST. Brains were next washed 4 times — each wash 15 minutes at RT in 0.5% PBST — and then rinsed in PBS for 15 minutes at RT. Brains were then mounted on PLL-coated coverslips in PBS, quickly dipped in MilliQ water, and dehydrated for 10 minutes in successively higher concentration of ethanol baths. Finally, the brains were soaked in 3 xylene baths for 5 minutes each, DPX was applied, the coverslip was mounted on a slide and was left in the hood for at least 48 hours before imaging. A detailed list of antibodies used in each figure is in Table 3-S1. The primary antibodies used for this study are: Goat anti-GFP (Rockland #600-101-215, 1:1000), Guinea Pig anti-FruM (Gift from Michael Perry, UCSD, 1:100) (Wohl et al., 2020), Guinea Pig anti-RFP (Gift from Susan Brenner-Morton, Columbia University, 1:10,000), Mouse anti-CD2 (Bio-Rad #MCA154GA, 1:100), Rabbit anti-RFP (Abcam #ab62341, 1:500 – only used in Figure 4-S4), Rat anti-HA (Roche, 11867423001; 1:100). It is important to note that the Goat anti-GFP antibody works much better for second-order staining of all LexAop-GFP containing flies than any of the other GFP antibodies we tried and much better than any of the V5 epitope tag antibodies we tried (in LexAop-GFP, the GFP has a V5 epitope tag attached). Secondary antibodies were spun down at 15,000 RPM for 15 minutes before adding to blocking buffer. The secondary antibodies used for this study are: Donkey anti-goat 488 (ThermoFisher, #A32814), Donkey anti-Guinea Pig Cy3 550 (Jackson Immuno, #706-165-148), Donkey anti-Guinea Pig 647 (EMD Millipore, #AP193SA6), Donkey anti-mouse 647 (ThermoFisher, #A31571), Donkey anti-Rabbit 555 (ThermoFisher, #A32794), Donkey anti-Rat 555 (SouthernBiotech, #6430-32).

### Thermogenetic silencing experiments

Thermogenetic silencing courtship experiments were performed in a custom behavior chamber that was based on the previously described Flybowl^57^. Flies that were assayed for our shibire^ts^ experiments were analyzed at either 21°C or 31°C by adjusting the temperature in our behavior chamber. Flies that were analyzed as part of these experiments were placed in 21°C temperature- and humidity-controlled incubators upon eclosion. Flies in the 31°C groups were placed in the chamber for at least 30 minutes before being assayed to allow flies to acclimate to the temperature increase. Flies were analyzed at age 5-7 days old. Each trial consisted of an experimental male and a w^11^^18^ male fly. w^11^^18^ males used as the courtship targets had a single wing clipped to allow differentiation between experimental fly and w^11^^18^ fly. Assays were performed under white light (∼45 μW/cm2) with a polarizing filter. Courtship videos were manually annotated for frames when unilateral wing extensions occurred. Resulting data was then analyzed using custom R scripts. Code is available upon request.

### Microscopy and Image Analysis

All images were taken on a confocal microscope (Zeiss, LSM800) with ZEN Blue software (version 2.3) using auto-Z brightness correction when appropriate, to maintain homogenous signal in the Z plane. Maximum intensity Z-stacks were created in ZEN and ultimately exported as TIFF files. All figures were generated in Adobe Illustrator (version 26.0.1).

### Electron Microscopy Circuit Reconstructions

*ds*-Tango EM simulations were performed using the natverse suite of R packages^75^. The first order populations were queried using the “neuprint_search” function. The “neuprint_simple_connectivity” function was used to retrieve lists of postsynaptic partners that were annotated by the number of synapses between each neuron. We thresholded first-to-second and second-to-third synaptic partnerships using the total number of synapses each neuron received from the first order and second order populations, respectively. We used a threshold of 30 for first-to-second order connections and 50 for second-to-third connections. The “neuprint_read_skeletons” function was used to retrieve EM skeletonizations and the “mirror_brain” function was used to mirror skeletons between right and left hemispheres.

## Acknowledgments

We would like to thank Dr. Alexander Fleischmann and the members of the Barnea Laboratory for critical reading of the manuscript. We acknowledge Susan Brenner-Morton for sharing reagents and John Murphy from the Brown BioMed Machine Shop for aiding in the construction of our behavior chamber. This work was supported by NIH/NIDCD 5R01DC017146 (G.B.), Brown University Carney Institute for Brain Science, Suna Kıraç Fund for Brain Science (D.S.), Brown University Carney Institute for Brain Science, Graduate Award in Brain Science (D.S.) and NIH/NIDCD award F31DC019540 (A.M.C.). Stocks obtained from the Bloomington *Drosophila* Stock Center (NIH P40OD018537) were used in this study.

## Author contributions

J.D.F., A.M.C., A.S., and G.B. conceptualized the study. J.D.F., A.M.C., A.S., and G.B. devised the methodology. J.D.F, A.M.C., A.S., S.M-M., N.J.S., N.V., S.M., A.V., A.H.W., R.A.M., A.M.O., D.S., B.N., P.I., A.K., C.C-F., G.G.H., and M.T contributed to the investigation and visualization of the results. J.D.F., A.M.C., A.S., and G.B. administered the project. J.D.F., A.M.C., A.S., and G.B. wrote the manuscript. The funding was obtained by, and the project was supervised by G.B.

## Additional information

All new fly strains will be deposited to Bloomington *Drosophila* Stock Center. Correspondence and requests for materials should be addressed to G.B.

