## Supplemental Table and Figures for "Convergent olfactory circuits for courtship in *Drosophila* revealed by *ds*-Tango"

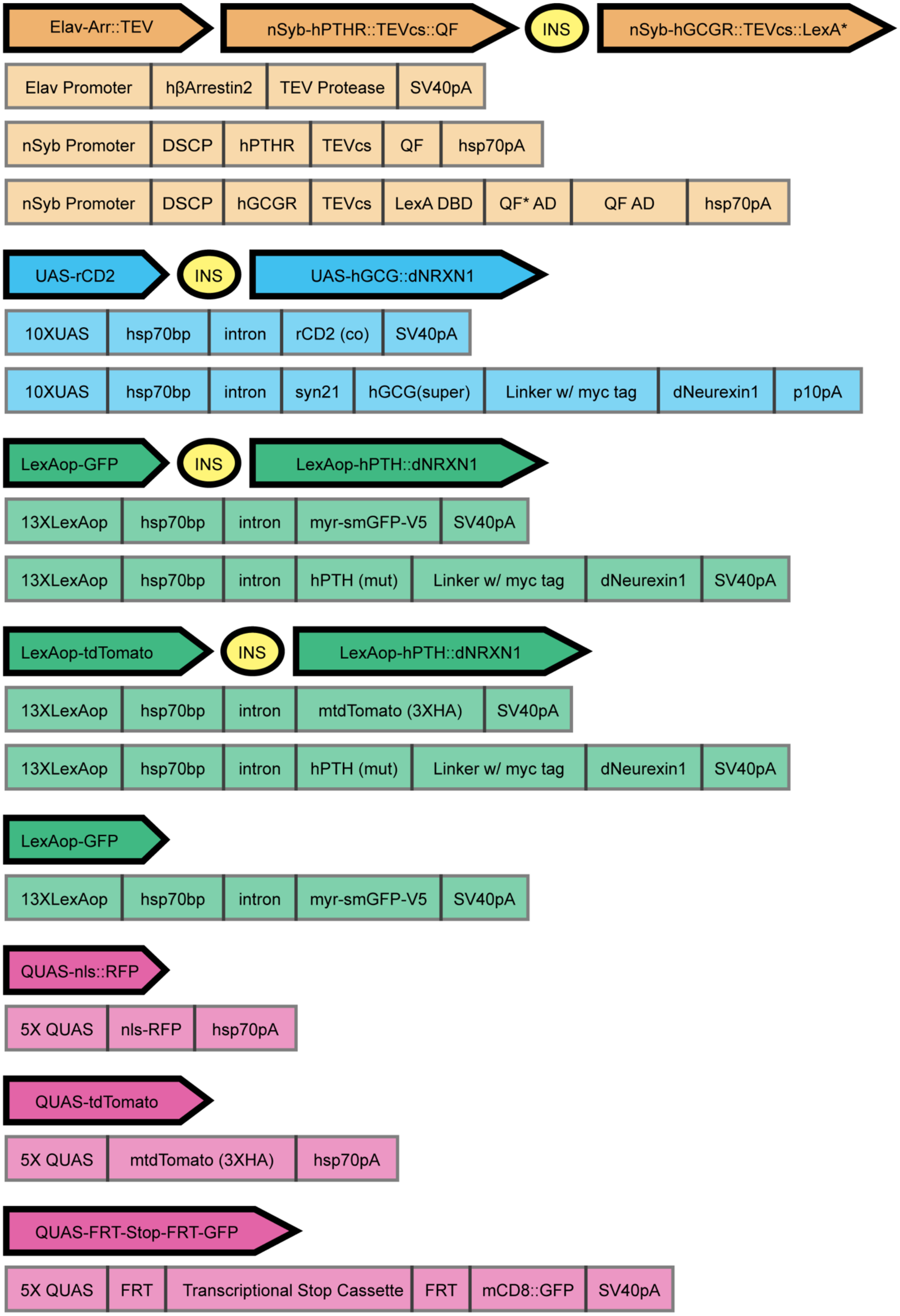
Elav-Arr::TEV
nSyb-hPTHR::TEVcs::QF
INS
nSyb-hGCGR::TEVcs::LexA*
Elav Promoter
hβArrestin2
TEV Protease
SV40pA
nSyb Promoter
DSCP
hPTHR
TEVcs
QF
hsp70pA
nSyb Promoter
DSCP
hGCGR
TEVcs
LexA DBD
QF* AD
QF AD
hsp70pA
UAS-rCD2
INS
UAS-hGCG::dNRXN1
10XUAS
hsp70bp
intron
rCD2 (co)
SV40pA
10XUAS
hsp70bp
intron
syn21
hGCG(super)
Linker w/ myc tag
dNeurexin1
p10pA
LexAop-GFP
INS
LexAop-hPTH::dNRXN1
13XLexAop
hsp70bp
intron
myr-smGFP-V5
SV40pA
13XLexAop
hsp70bp
intron
hPTH (mut)
Linker w/ myc tag
dNeurexin1
SV40pA
LexAop-tdTomato
INS
LexAop-hPTH::dNRXN1
13XLexAop
hsp70bp
intron
mtdTomato (3XHA)
SV40pA
13XLexAop
hsp70bp
intron
hPTH (mut)
Linker w/ myc tag
dNeurexin1
SV40pA
LexAop-GFP
13XLexAop
hsp70bp
intron
myr-smGFP-V5
SV40pA
QUAS-nls::RFP
5X QUAS
nls-RFP
hsp70pA
QUAS-tdTomato
5X QUAS
mtdTomato (3XHA)
hsp70pA
QUAS-FRT-Stop-FRT-GFP
5X QUAS
FRT
Transcriptional Stop Cassette
FRT
mCD8::GFP
SV40pA

Figure S1. Detailed genetic components of *ds*-Tango flies

Detailed descriptions of the genetic components of *ds*-Tango flies used in this study. The presence of a gypsy insulator (INS) in an allele is indicated with a circle (yellow). One allele of panneuronally expressed *ds*-Tango components (brown). One allele of Gal4-dependent *ds*-Tango components (blue). Three separate alleles of LexA*-dependent *ds*-Tango components (green). Three separate alleles of QF-dependent *ds*-Tango components (magenta). Bolded components that are horizontal to one another indicate they were inserted as a single allele. Below every individual allele, further detail on each of the genetic components (light colored rectangles) making up the allele is shown. See methods and supplemental text for further description of genetic components, including insertion sites. Elav: *Drosophila melanogaster* panneuronal promoter; pA: polyadenylation signal; nSyb: *Drosophila melanogaster* panneuronal promoter; DSCP: *Drosophila* Synthetic Core Promoter; hGCGR: human Glucagon Receptor; TEVcs: cleavage site for N1a protease from the Tobacco Etch Virus; UAS: Upstream Activating Sequence for Gal4; hGCG: human Glucagon analogue with enhanced receptor binding; GFP: Green Fluorescent Protein; QUAS: Upstream Activating Sequence for QF; INS: gypsy insulator; hPTH: human Parathyroid Hormone; hPTHR: human Parathyroid Hormone Receptor

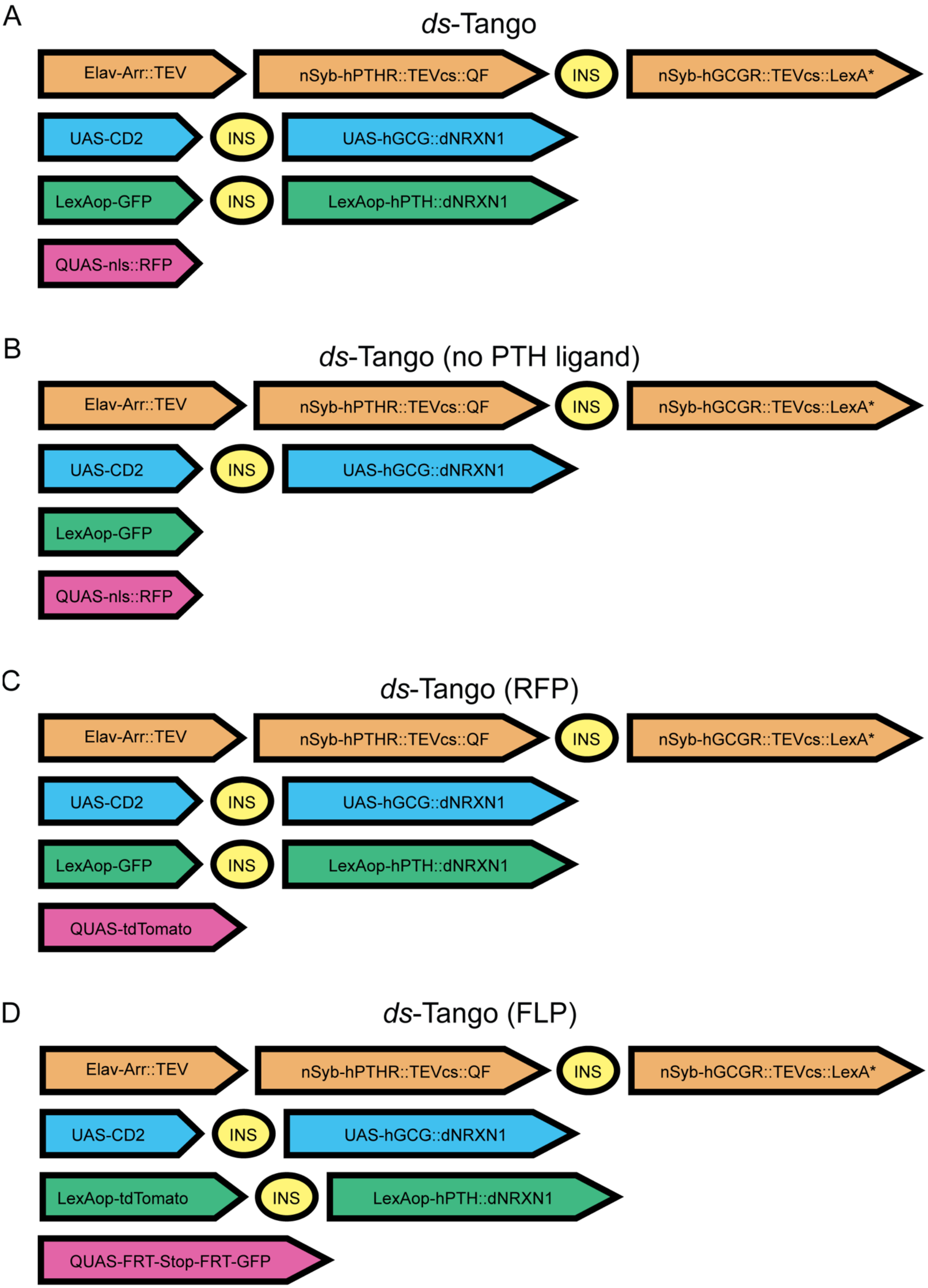
A
*ds*-Tango
Elav-Arr::TEV
nSyb-hPTHR::TEVcs::QF
INS
nSyb-hGCGR::TEVcs::LexA*
UAS-CD2
INS
UAS-hGCG::dNRXN1
LexAop-GFP
INS
LexAop-hPTH::dNRXN1
QUAS-nls::RFP
B
*ds*-Tango (no PTH ligand)
Elav-Arr::TEV
nSyb-hPTHR::TEVcs::QF
INS
nSyb-hGCGR::TEVcs::LexA*
UAS-CD2
INS
UAS-hGCG::dNRXN1
LexAop-GFP
QUAS-nls::RFP
C
*ds*-Tango (RFP)
Elav-Arr::TEV
nSyb-hPTHR::TEVcs::QF
INS
nSyb-hGCGR::TEVcs::LexA*
UAS-CD2
INS
UAS-hGCG::dNRXN1
LexAop-GFP
INS
LexAop-hPTH::dNRXN1
QUAS-tdTomato
D
*ds*-Tango (FLP)
Elav-Arr::TEV
nSyb-hPTHR::TEVcs::QF
INS
nSyb-hGCGR::TEVcs::LexA*
UAS-CD2
INS
UAS-hGCG::dNRXN1
LexAop-tdTomato
INS
LexAop-hPTH::dNRXN1
QUAS-FRT-Stop-FRT-GFP

Figure S2. Broad genetic components in different configurations of *ds*-Tango

(A) The genetic components in “*ds*-Tango” flies.

(B) The genetic components in “*ds*-Tango (no PTH ligand)” flies. When compared to (A), note the lack of both a gypsy insulator (INS) (yellow) and the LexAop-hPTH::dNRXN1 sequence (green).

(C) The genetic components in “*ds*-Tango (RFP)” flies. Note these flies have a different, non-nuclear localized, disynaptic reporter (magenta) compared to the flies in (A).

(D) The genetic components in “*ds*-Tango (FLP)” flies. When compared to (A), note the different LexAop reporter, LexAop-tdTomato (green), and the different FLP-dependent third-order reporter (magenta).

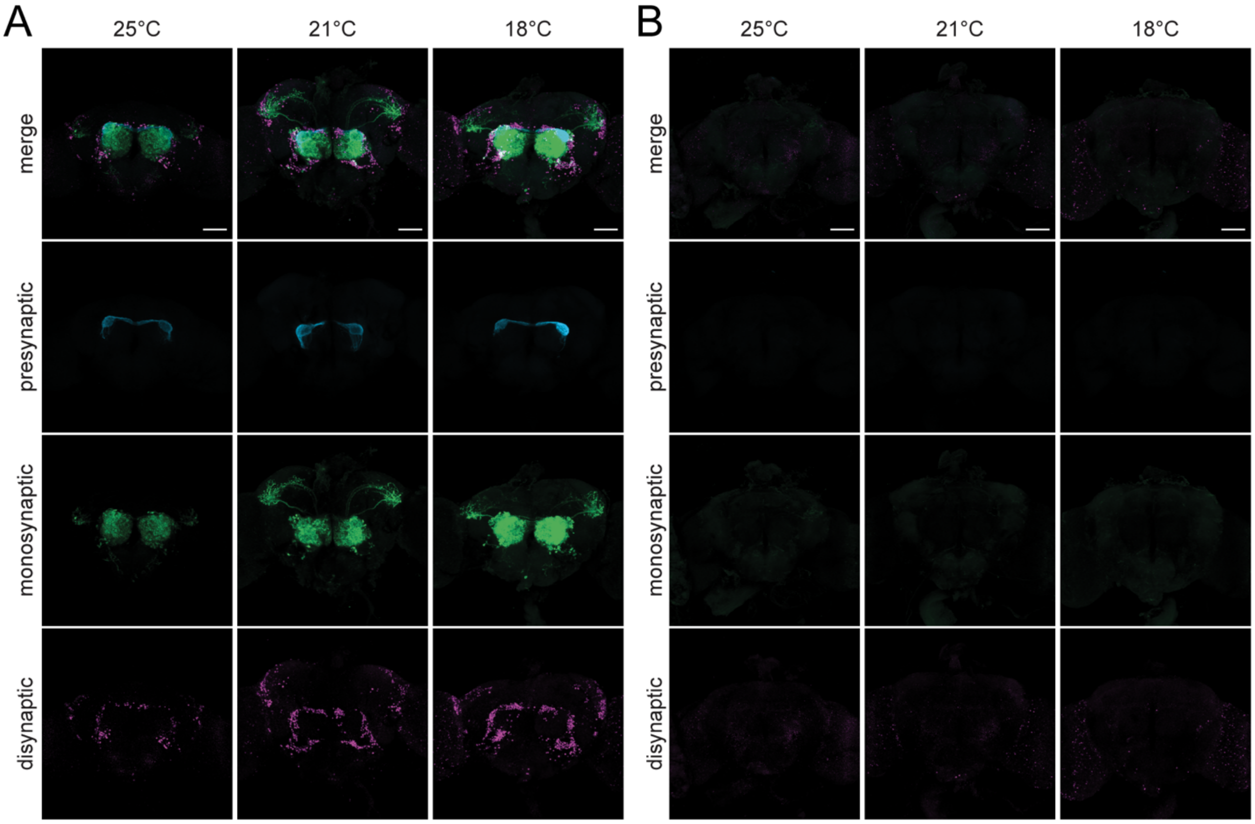


Figure S3. The effect of temperature on *ds*-Tango

(A) *ds*-Tango signal when initiated from Or67d-OSNs in flies with identical experimental parameters and image settings but crossed and raised at different temperatures (labeled at top of figure). Or67d-OSNs (cyan), their monosynaptic partners LNs and OPNs (green), and the nuclei of their disynaptic connections (magenta) are shown.

(B) No Gal4 control *ds*-Tango signal from flies with identical experimental parameters and image settings but crossed and raised at different temperatures. Presynaptic background signal (cyan), monosynaptic background signal (green), and the nuclei of the neurons where disynaptic background signal is present (magenta) are shown. Individual channels are shown below each merged channel image. Each image is a maximum intensity Z-stack projection of a whole-mount brain. Scale bars, 50µm.

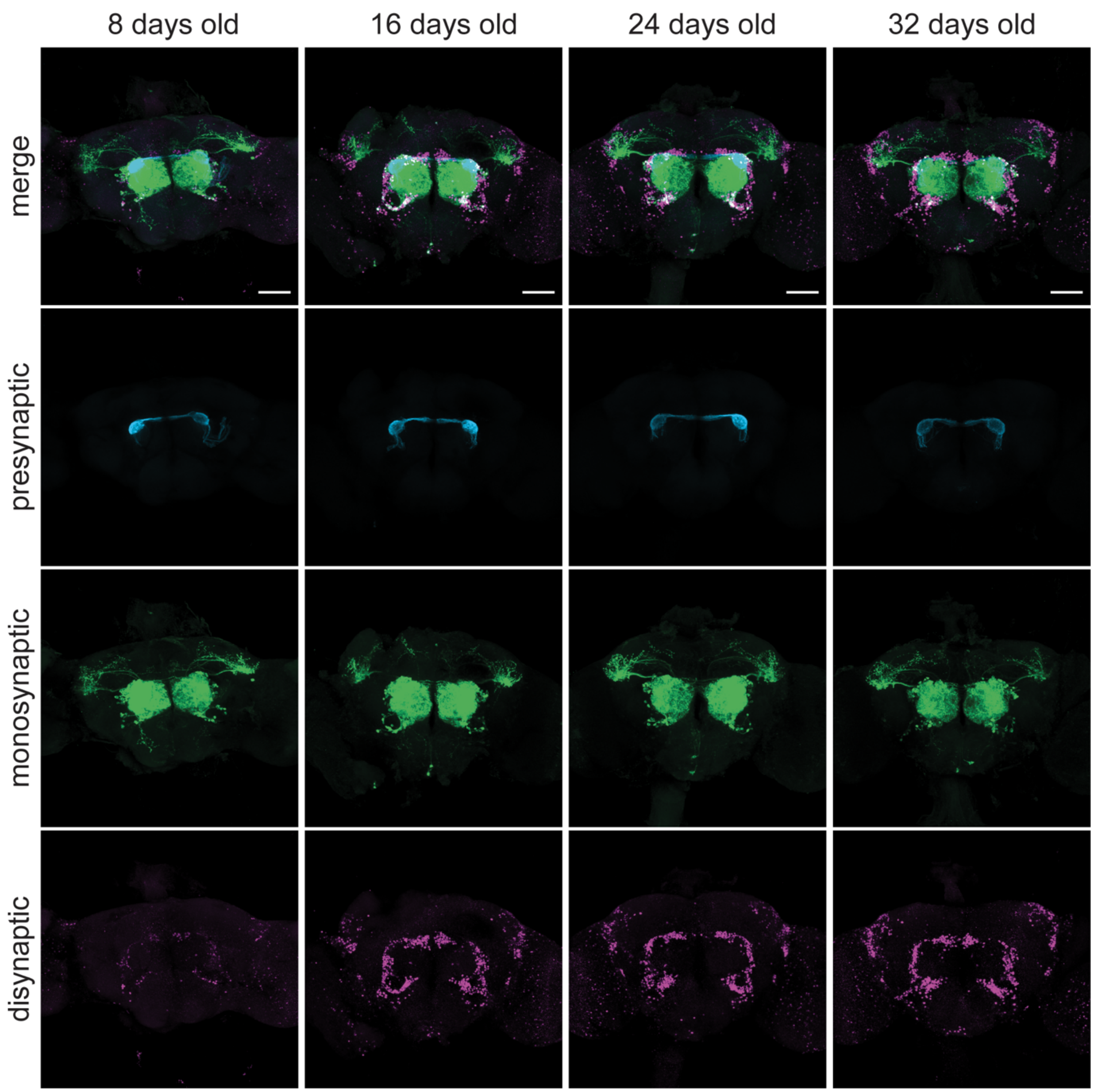


Figure S4. Age Dependence of the *ds*-Tango Signal

*ds*-Tango signal when initiated from Or67d-OSNs in flies with identical experimental parameters and image settings but dissected at different ages (labeled at top of figure). Or67d-OSNs (cyan), their monosynaptic partners LNs and OPNs (green), and the nuclei of their disynaptic connections (magenta) are shown. Individual channels are shown below each merged channel image. Each image is a maximum intensity Z-stack projection of a whole-mount brain. Scale bars, 50µm.

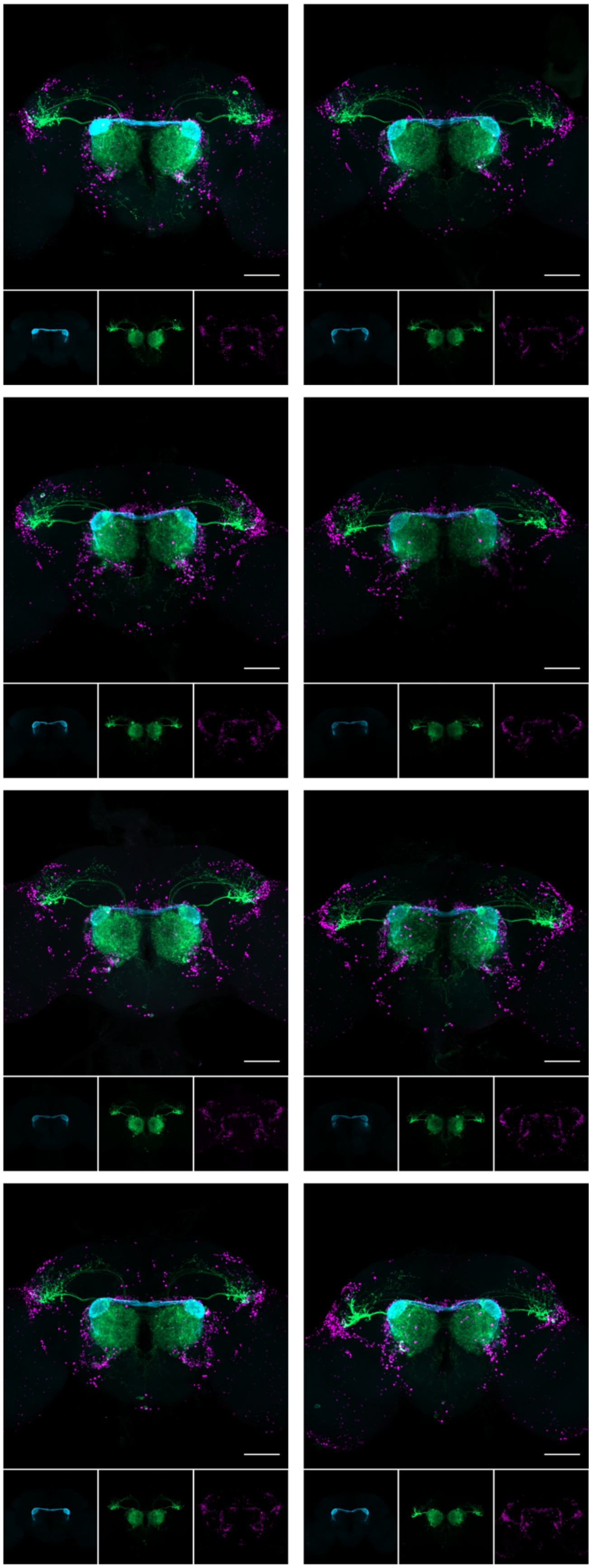

Figure S5. Consistency of the *ds*-Tango Signal

*ds*-Tango signal when initiated from Or67d-OSNs in eight different male brains with the same experimental parameters and image settings. Or67d-OSNs (cyan), their monosynaptic partners LNs and OPNs (green), and the nuclei of their disynaptic connections (magenta) are shown. Below each image of a brain, the individual channels for that image are displayed. Each image is a maximum intensity Z-stack projection of a whole-mount brain. Scale bars, 50µm.

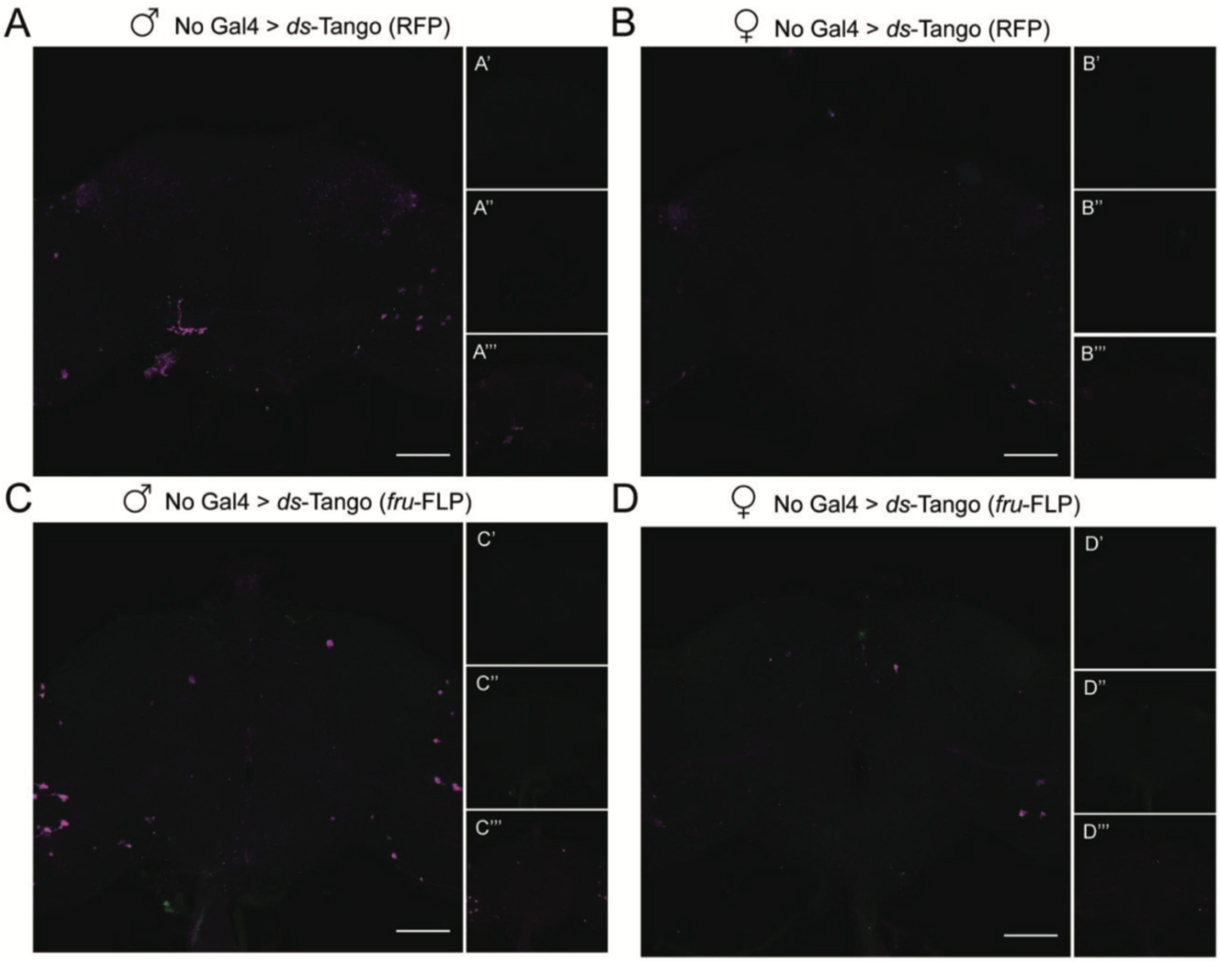


Figure S6. Minimal background signal is observed in different configurations of *ds*-Tango

A control experiment driving *ds*-Tango with a membrane-localized third-order reporter (RFP) and no Gal4 driver in males (A) and in females (B). Presynaptic background signal (cyan in A', B'), monosynaptic background signal (green in A'', B''), and disynaptic background signal (magenta in A''', B''') are shown.

A control experiment driving *ds*-Tango with a fru-FLP dependent reporter and no Gal4 driver in males (C) and in females (D). Presynaptic background signal (cyan shown in C', D'), monosynaptic background signal (green shown in C'', D''), and fru-FLP+ neurons where disynaptic background signal is present (magenta shown in C''', D''') are shown. Each image is a maximum intensity Z-stack projection of a whole-mount brain. Scale bars, 50µm.

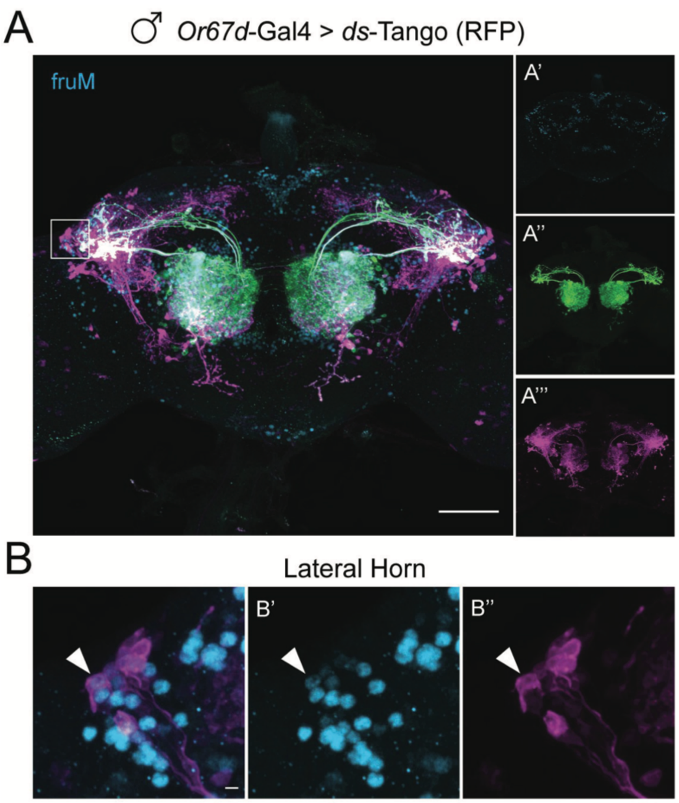


Figure S7. Some disynaptic connections revealed by *ds*-Tango when initiated from Or67d-OSNs are fruM+

(A) Driving the *ds*-Tango configuration containing a membrane-localized third-order reporter (RFP) with *Or67d*-Gal4 in males. fruM antibody staining (cyan, in A'), the monosynaptic partners LNs and OPNs (green, in A''), and the disynaptic connections (magenta, in A''') are shown.

(B) A higher magnification image of the gray inset in (A) highlighting the left LH reveals colabeling of fruM (cyan, channel show in B') and some third-order LC neurons (magenta, shown in B''). Maximum intensity full Z-stack projection (A) and partial Z-stack projection (B) of whole-mount brains are shown. Scale bars, 50µm.

A

B

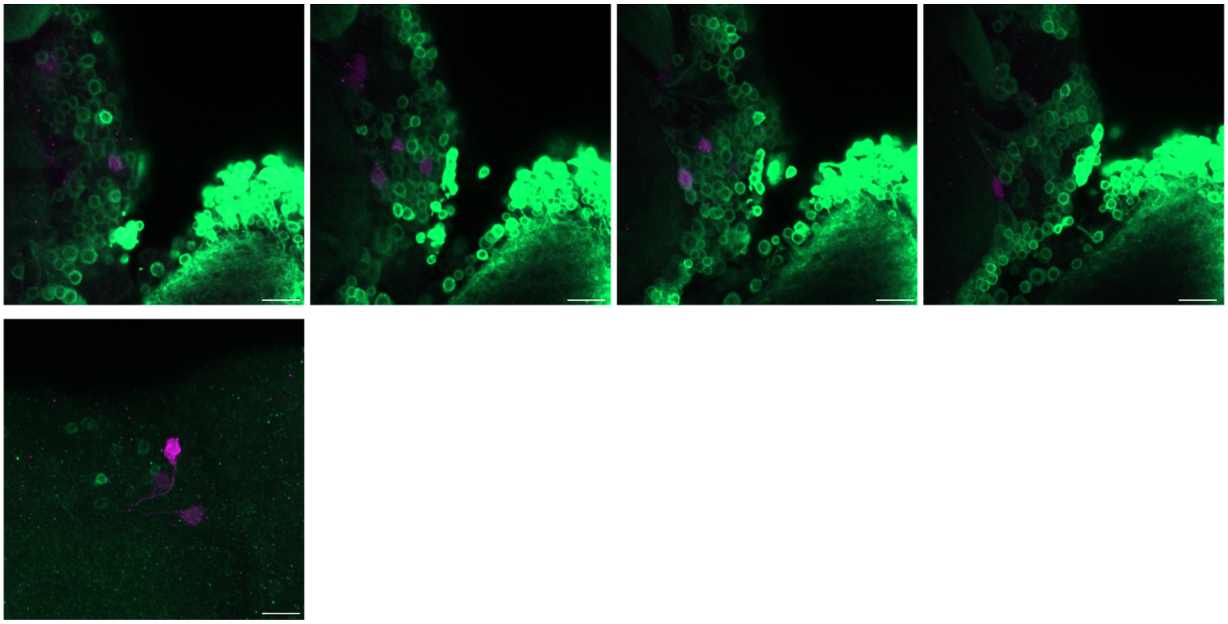

Figure S8. Almost all Maiandros neurons are cholinergic in males

(A) Subsets of the Z-stacks of the whole brain showing the Maiandros neurons (magenta) and GFP expressed under the control of the ChAT promoter (green). Almost all Maiandros neurons express GFP.

(B) Subset of the Z-stacks of the whole brain showing the Maiandros neurons (magenta) and GFP expressed under the control of the GAD promoter (green). No Maiandros neuron expresses GFP.

| Figure | Genotype | Antibodies | Sex | Age (weeks) | Temp (°C) |
|---|---|---|---|---|---|
| 1A | QUAS-mtdTomato-HA, UAS-myrGFP; *trans*-Tango/+; *Or67d*- Gal4/+ | Cyan-GFP, Red-RFP, Grey-nc82 | M | 3-4 | 18 |
| 1B | QUAS-mtdTomato-HA, UAS-myrGFP; *trans*-Tango/+; *Or47b*- Gal4/+ | Cyan-GFP, Red-RFP, Grey-nc82 | M | 3-4 | 18 |
| 1C | QUAS-mtdTomato-HA, UAS-myrGFP; *trans*-Tango/+; *Ir84a*- Gal4/+ | Cyan-GFP, Red-RFP, Grey-nc82 | M | 3-4 | 18 |
| 2B | QUAS-nls::RFP; Elav-Arr:TEV nSyb-PTHR::TEVcs::QF INS nSyb-GCGR::TEVcs::LexA*/+; UAS-CD2 INS UAS-GCG::dNRX1, LexAop-GFP INS LexAop-PTH::dNRX1/*Orco*-Gal4 | Cyan-CD2, Green-GFP, Magenta-RFP | F | 3-4 | 21 |
| 2C | QUAS-nls::RFP; Elav-Arr:TEV nSyb-PTHR::TEVcs::QF INS nSyb-GCGR::TEVcs::LexA*/+; UAS-CD2 INS UAS-GCG::dNRX1, LexAop-GFP INS LexAop-PTH::dNRX1/+ | Cyan-CD2, Green-GFP, Magenta-RFP | F | 3-4 | 21 |
| 2E | QUAS-nls::RFP; Elav-Arr:TEV nSyb-PTHR::TEVcs::QF INS nSyb-GCGR::TEVcs::LexA*/+; UAS-CD2 INS UAS-GCG::dNRX1, LexAop-GFP/Orco-Gal4 | Cyan-CD2, Green-GFP, Magenta-RFP | F | 3-4 | 21 |
| 3B, 3C | QUAS-nls::RFP; Elav-Arr:TEV nSyb-PTHR::TEVcs::QF INS nSyb-GCGR::TEVcs::LexA*/+; UAS-CD2 INS UAS-GCG::dNRX1, LexAop-GFP INS LexAop-PTH::dNRX1/*Or67d*-Gal4 | Cyan-CD2, Green-GFP, Magenta-RFP | F | 3-4 | 21 |
| 3E, 3F | QUAS-nls::RFP; Elav-Arr:TEV nSyb-PTHR::TEVcs::QF INS nSyb-GCGR::TEVcs::LexA*/+; UAS-CD2 INS UAS-GCG::dNRX1, LexAop-GFP INS LexAop-PTH::dNRX1/*Or42a*-Gal4 | Cyan-CD2, Green-GFP, Magenta-RFP | F | 3-4 | 21 |
| 4A | QUAS-tdTomato; Elav-Arr:TEV nSyb-PTHR::TEVcs::QF INS nSyb-GCGR::TEVcs::LexA*/+; UAS-CD2 INS UAS-GCG::dNRX1, LexAop-GFP INS LexAop-PTH::dNRX1/*Or67*d-Gal4 | Cyan-CD2, Green-GFP, Magenta-HA | M | 3-4 | 25 |
| 4B, 4C | w; Elav-Arr:TEV nSyb-PTHR::TEVcs::QF INS nSyb-GCGR::TEVcs::LexA*/QUAS-FRT- STOP-FRT-GFP; UAS-CD2 INS UAS-GCG::dNRX1, LexAop-tdTomato INS LexAop-PTH::dNRX1/*Or67d*-Gal4, *fru*-FLP | Cyan-CD2, Green-RFP, Magenta-GFP | M | 4 | 18 |
| 4D | QUAS-tdTomato/+; Elav-Arr:TEV nSyb-PTHR::TEVcs::QF INS nSyb-GCGR::TEVcs::LexA*/+; UAS-CD2 INS UAS-GCG::dNRX1, LexAop-GFP INS LexAop-PTH::dNRX1/*Or67*d-Gal4 | Cyan-CD2, Green-GFP, Magenta-HA | F | 3-4 | 25 |

| | | | | | |
|---|---|---|---|---|---|
| 4E, 4F | w; Elav-Arr:TEV nSyb-PTHR::TEVcs::QF INS nSyb-GCGR::TEVcs::LexA*/QUAS-FRT- STOP-FRT-GFP; UAS-CD2 INS UAS-GCG::dNRX1, LexAop-tdTomato INS LexAop-PTH::dNRX1/*Or67d*-Gal4, *fru*-FLP | Cyan-CD2, Green-RFP, Magenta-GFP | F | 4 | 18 |
| 5A | QUAS-tdTomato; Elav-Arr:TEV nSyb-PTHR::TEVcs::QF INS nSyb-GCGR::TEVcs::LexA*/*Or47b*-Gal4; UAS-CD2 INS UAS-GCG::dNRX1, LexAop-GFP INS LexAop-PTH::dNRX1/+ | Cyan-CD2, Green-RFP, Magenta-GFP | M | 3-4 | 21 |
| 5B | w; Elav-Arr:TEV nSyb-PTHR::TEVcs::QF INS nSyb-GCGR::TEVcs::LexA*/QUAS-FRT- STOP-FRT-GFP/*Or47b*-Gal4; UAS-CD2 INS UAS-GCG::dNRX1, LexAop-tdTomato INS LexAop-PTH::dNRX1/ *fru*-FLP | Cyan-CD2, Green-RFP, Magenta-GFP | M | 3-4 | 21 |
| 5C | QUAS-tdTomato; Elav-Arr:TEV nSyb-PTHR::TEVcs::QF INS nSyb-GCGR::TEVcs::LexA*/*Ir84a*-Gal4; UAS-CD2 INS UAS-GCG::dNRX1, LexAop-GFP INS LexAop-PTH::dNRX1/+ | Cyan-CD2, Green-RFP, Magenta-GFP | M | 3-4 | 21 |
| 5D | w; Elav-Arr:TEV nSyb-PTHR::TEVcs::QF INS nSyb-GCGR::TEVcs::LexA*/QUAS-FRT- STOP-FRT-GFP/*Ir84a*-Gal4; UAS-CD2 INS UAS-GCG::dNRX1, LexAop-tdTomato INS LexAop-PTH::dNRX1/ *fru*-FLP | Cyan-CD2, Green-RFP, Magenta-GFP | M | 3-4 | 21 |
| 6A | +;VT060731-p65ADZp/+; VT006486-ZpGDBD /UAS-GFP | Green-GFP Grey-nc82 | M | 1 | 25 |
| 6B | +;VT060731-p65ADZp/+; VT006486-ZpGDBD /UAS-Syt::GFP,UAS-DenMark | Green-GFP Magenta-DenMark Grey-nc82 | M | 1 | 25 |
| 6C, 6D, 6E | +;VT060731-p65ADZp/+; VT006486-ZpGDBD /UAS-Shi$^{TS}$<br>+;+/+;+/ Shi$^{TS}$ | NA | M | 1 | 21 |
| S3A | QUAS-nls::RFP; Elav-Arr:TEV nSyb-PTHR::TEVcs::QF INS nSyb-GCGR::TEVcs::LexA*/+; UAS-CD2 INS UAS-GCG::dNRX1, LexAop-GFP INS LexAop-PTH::dNRX1/*Or67d*-Gal4 | Cyan-CD2, Green-GFP, Magenta-RFP | M | 3-4 | N/A |
| S3B | QUAS-nls::RFP; Elav-Arr:TEV nSyb-PTHR::TEVcs::QF INS nSyb-GCGR::TEVcs::LexA*/+; UAS-CD2 INS UAS-GCG::dNRX1, LexAop-GFP INS LexAop-PTH::dNRX1/+ | Cyan-CD2, Green-GFP, Magenta-RFP | M | 3-4 | N/A |
| S4 | QUAS-nls::RFP; Elav-Arr:TEV nSyb-PTHR::TEVcs::QF INS nSyb-GCGR::TEVcs::LexA*/+; UAS-CD2 INS UAS-GCG::dNRX1, LexAop-GFP INS LexAop-PTH::dNRX1/*Or67d*-Gal4 | Cyan-CD2, Green-GFP, Magenta-RFP | M | N/A | 18 |

| | | | | | |
|---|---|---|---|---|---|
| S5 | QUAS-nls::RFP; Elav-Arr:TEV nSyb-PTHR::TEVcs::QF INS nSyb-GCGR::TEVcs::LexA*/+; UAS-CD2 INS UAS-GCG::dNRX1, LexAop-GFP INS LexAop-PTH::dNRX1/*Or67d*-Gal4 | Cyan-CD2, Green-GFP, Magenta-RFP | M | 3-4 | 21 |
| S6A | QUAS-tdTomato; Elav-Arr:TEV nSyb-PTHR::TEVcs::QF INS nSyb-GCGR::TEVcs::LexA*/+; UAS-CD2 INS UAS-GCG::dNRX1, LexAop-GFP INS LexAop-PTH::dNRX1/+ | Cyan-CD2, Green-GFP, Magenta-HA | M | 3-4 | 25 |
| S6B | QUAS-tdTomato/+; Elav-Arr:TEV nSyb-PTHR::TEVcs::QF INS nSyb-GCGR::TEVcs::LexA*/+; UAS-CD2 INS UAS-GCG::dNRX1, LexAop-GFP INS LexAop-PTH::dNRX1/+ | Cyan-CD2, Green-GFP, Magenta-HA | F | 3-4 | 25 |
| S6C | w; Elav-Arr:TEV nSyb-PTHR::TEVcs::QF INS nSyb-GCGR::TEVcs::LexA*/QUAS-FRT- STOP-FRT-GFP; UAS-CD2 INS UAS-GCG::dNRX1, LexAop-tdTomato INS LexAop-PTH::dNRX1/ *fru*-FLP | Cyan-CD2, Green-RFP, Magenta-GFP | M | 4 | 18 |
| S6D | w; Elav-Arr:TEV nSyb-PTHR::TEVcs::QF INS nSyb-GCGR::TEVcs::LexA*/QUAS-FRT- STOP-FRT-GFP; UAS-CD2 INS UAS-GCG::dNRX1, LexAop-tdTomato INS LexAop-PTH::dNRX1/*fru*-FLP | Cyan-CD2, Green-RFP, Magenta-GFP | F | 4 | 18 |
| S7 | QUAS-tdTomato; Elav-Arr:TEV nSyb-PTHR::TEVcs::QF INS nSyb-GCGR::TEVcs::LexA*/+; UAS-CD2 INS UAS-GCG::dNRX1, LexAop-GFP INS LexAop-PTH::dNRX1/*Or67*d-Gal4 | Cyan-fruM, Green-GFP, Magenta-HA | M | 3-4 | 21 |
| S8A | UAS-mCD8-RFP, LexAop-mCD8-GFP;VT060731-p65ADZp/+; VT006486-ZpGDBD/ChAT-P2A-LexA | Green-GFP Magenta-RFP | M | 1 | 25 |
| S8B | UAS-mCD8-RFP, LexAop-mCD8-GFP;VT060731-p65ADZp/+; VT006486-ZpGDBD/GAD-P2A-LexA | Green-GFP Magenta-RFP | M | 1 | 25 |

Table S1. Fly genotypes used and experimental details for each figure
Old
