## Supplemental Text for "Convergent olfactory circuits for courtship in *Drosophila* revealed by *ds*-Tango"

*ds*-Tango is based on the established *trans*-Tango system for circuit mapping[1], but instead enables differential access into neurons that are mono- and disynaptic to the starting population. Conceptually, *ds*-Tango relies on two *trans*-Tango systems that are configured in series. Both *ds*- and *trans*-Tango are based on the Tango assay, a cell culture assay which translates cell surface activation of a receptor, by a ligand, into stable reporter gene expression in the cell[2]. There are 3 components to the Tango assay, a membrane-bound GPCR that is tethered to a transcription factor via a tobacco etch virus (TEV) protease cleavage site, a β-Arrestin-TEV fusion protein, and a transcription factor responsive element in the nucleus. In the absence of the ligand, the transcription factor stays tethered to the membrane and no reporter gene is expressed. However, upon ligand binding, the receptor is activated, recruiting the cytosolic signaling molecule β-Arrestin along with the fused TEV protease. The protease cuts the cleavage site between the receptor and the transcription factor, releasing the transcription factor, which is then free to translocate to the nucleus and drive transcription of a reporter gene. In *trans*-Tango, the genetically encoded monosynaptic anterograde tracing technique we previously developed, all the components of the glucagon receptor (GCGR) version of Tango were introduced into all neurons of the fly[1]. We then expressed a tethered glucagon (GCG) ligand in the presynaptic terminals of genetically defined subsets of neurons. The presynaptic ligand activates the Tango GCGR on the postsynaptic neurons, setting off the Tango signaling cascade exclusively in postsynaptic cells; thereby enabling genetic access to postsynaptic neurons of the defined circuit. Several modifications were made to the monosynaptic tracer *trans*-Tango to enable disynaptic tracing using *ds*-Tango.

First, we developed a second *trans*-Tango ligand-receptor pair that acts via a membrane tethered version of the ligand, is exogenous to *Drosophila*, exhibits a high signal-to-noise ratio, and does not cross react with our first *trans*-Tango ligand-receptor pair, GCG-GCGR. We first screened through other class B1 GPCRs that have been shown to be activated by membrane-tethered ligands[3]. The human parathyroid hormone (PTH) and parathyroid hormone receptor (PTHR) were identified and optimized in the fly, *in vivo,* to most accurately recapitulate the signal that the GCG/GCGR version of *trans*-Tango initially produced (data not shown). Importantly, both human PTH and PTHR are exogenous to *Drosophila*. Some amino acids were deleted from the human parathyroid hormone ligand to improve signal-to-noise with *trans*-Tango, yielding a mutated parathyroid hormone (PTH) that we used for all experiments in this study. The PTH ligand was further shown to not cross react with the GCGR, and the GCG ligand was shown to not cross react with the PTHR, validating that two separate *trans*-Tango ligand-receptor pairs that can act in a single animal had been developed.

Next, we had to develop a second transcription factor that could be fused to the Tango version of either receptor, such that the activation of each receptor-transcription factor fusion would generate a distinct transcriptional readout. The LexA/LexAop binary system was appealing for the potential development of a second transcription factor because we were already using the binary systems Gal4/UAS as our genetic entry in *trans*-Tango and QF/QUAS as our genetic exit[1,4-6]. Further, all three binary systems have been shown to successfully work in concert in the fly nervous system without cross-reacting[7] (Riabinina and Potter 2016). To optimize the LexA transcription factor for use in the context of *trans*-

Tango, we had to make two modifications. First, we had to replace the VP16 or p65 activation domains (AD) of LexA with the QF AD to observe any *trans*-Tango postsynaptic signal. Secondly, we had to insert a flexible linker domain between the LexA DNA-binding domain (DBD) and the QF AD of the new transcription factor in order to improve the signal, such that it recapitulates the original QF *trans*-Tango postsynaptic signal. We developed two linkers: one that exhibited stronger *trans*-Tango signal, with flexible Glycine-Serine repeats and another that exhibited weaker *trans*-Tango signal, with part of the endogenous QF middle domain, termed QF* (from QFf in the original article)[7]. Ultimately, we settled on this weaker LexA DBD/QF*/QF AD hybrid transcription factor, LexA*, because it exhibited less background noise, and therefore a higher signal-to-noise ratio than the Glycine-Serine version.

Next, we modified both ligand fusion proteins from the original *trans*-Tango design by replacing the human ICAM sequence with the full-length *Drosophila* Neurexin1 (dNRXN1). We observed a higher expression of this version of the ligand fusion, perhaps because dNRXN1 is already codon-optimized for *Drosophila*. The use of this version of the ligand fusion yielded higher signal-to-noise ratio than the version used in the original *trans*-Tango.

Finally, we optimized the ligands and reporters under control of each transcription factor by placing two separate optimized DNA binding domains for each ligand and reporter separated by a gypsy-insulated spacer (Figures S1 and S2, and further details in Methods)[8]. The insulator provided for a higher level of expression of both ligand and reporter than if the construct lacked the gypsy sequence. Another key difference from the original *trans*-Tango is that in *ds*-Tango any two transcription factor responsive elements that were the same, were placed in the same insertion site in the *Drosophila* genome. For instance, in the original *trans*-Tango the UAS-GCG ligand construct was inserted into the attP40 locus on the second chromosome, while the UAS-GFP reporter construct was inserted into the Su(Hw)attP8 locus on the X chromosome. This led to some instances where GFP would be present where GCG ligand was not, and vice versa.

To stably introduce all the necessary *ds*-Tango genetic components into the fly, we generated four separate constructs that were independently incorporated into the fly genome using PhiC31-mediated integration. First, we expressed the Arr:TEV fusion protein, PTHR::TEVcs::QF fusion protein, and GCGR::TEVcs::LexA* fusion protein in all neurons of the fly using panneuronal promoters (Figure S1 and further details in Methods). These three fusion proteins were cloned into a single plasmid that was inserted into the attP40 insertion site of the fly. Second, we inserted a single plasmid containing both the UAS-CD2 reporter and UAS-GCG::dNRX ligand fusion construct into the attp2 insertion site on the third chromosome (Figure S1 and further details in Methods). Both the CD2 reporter and the GCG::dNRX ligand are expressed in a Gal4-dependent manner. Third, we inserted a single plasmid containing both the LexAop-GFP reporter and the LexAop-PTH-dNRX ligand fusion protein into the VK00027 insertion site on the third chromosome (Figure S1 and further details in Methods). Both the GFP reporter and the PTH::dNRX ligand are expressed in a LexA*-dependent manner. The fourth and final *ds*-Tango genetic component, a QUAS-nls::RFP reporter was integrated into the Su(Hw)attP8 insertion site on the X chromosome[9]. The nls::RFP reporter protein is expressed in a QF-dependent manner. The four *ds*-Tango genetic components were placed at different insertion sites to avoid transvection, and the insertion sites were selected based on their cumulative effects on signal-to-noise. Experimentally, flies bearing all of these alleles, termed *ds*-Tango flies, were crossed to Gal4 lines to enable discrete and restricted disynaptic tracing from the subsets of neurons that express Gal4 (a description of the four separate flies bearing different *ds*-Tango components used in this study are in Figure S2).

8. Pfeiffer, B.D., Truman, J.W., and Rubin, G.M. (2012). Using translational enhancers to increase transgene expression in Drosophila. Proc Natl Acad Sci U S A *109*, 6626-6631. 10.1073/pnas.1204520109.
9. Sorkac, A., Mosneanu, R.A., Crown, A.M., Savas, D., Okoro, A.M., Memis, E., Talay, M., and Barnea, G. (2023). retro-Tango enables versatile retrograde circuit tracing in Drosophila. Elife *12*. 10.7554/eLife.85041.
